# Improvement of sensory deficits in Fragile X mice by increasing cortical interneuron activity after the critical period

**DOI:** 10.1101/2022.05.17.492368

**Authors:** Nazim Kourdougli, Anand Suresh, Benjamin Liu, Pablo Juarez, Ashley Lin, David T. Chung, Anette Graven Sams, Michael Gandal, Verónica Martínez-Cerdeño, Dean V. Buonomano, Benjamin J. Hall, Cédric Mombereau, Carlos Portera-Cailliau

**Affiliations:** Department of Neurology, at the University of California Los Angeles (UCLA); Department of Neurobiology, at the University of California Los Angeles (UCLA); Department of Psychiatry of the David Geffen School of Medicine at the University of California Los Angeles (UCLA); Department of Pathology and Laboratory Medicine, UC Davis School of Medicine; MIND Institute for Pediatric Regenerative Medicine and Shriners Hospitals for Children, Sacramento, CA 95817, USA; Department of Psychology, UCLA; Lundbeck A/S H Ottiliavej 9, 2500 Copenhagen, Denmark

**Author notes:** Lead contact: Carlos Portera-Cailliau, MD, PhD, RNRC A-145, 710 Westwood Plaza, Los Angeles, CA 90095.

**Keywords:** In vivo calcium imaging, Autism spectrum disorders, Intellectual disability, Interneuron, Kv3.1, Medial ganglionic eminence, Parvalbumin, Ribotag, RNAseq, Somatosensory cortex, Transcriptomics, Two-photon

## Abstract

Changes in the function of inhibitory interneurons (INs) during cortical development could contribute to the pathophysiology of neurodevelopmental disorders. Using all-optical in vivo approaches in postnatal mouse somatosensory cortex (S1), we find that parvalbumin (PV) IN precursors are hypoactive and decoupled from excitatory neurons in *Fmr1^−/−^* mice, a model of Fragile X Syndrome (FXS). This leads to a loss of PV-INs in both mice and humans with FXS. Increasing the activity of future PV-INs in neonatal *Fmr1^−/−^* mice restores PV density and ameliorates transcriptional dysregulation in S1, but not circuit dysfunction. Critically, administering a novel allosteric modulator of Kv3.1 channels after the S1 critical period does rescue circuit dynamics and tactile defensiveness. Symptoms in FXS and related disorders could be mitigated by targeting PV-INs.

## INTRODUCTION

Neurodevelopmental disorders (NDDs) arise due to changes in developmental trajectories of neurons during the early stages of circuit assembly in the brain. Although symptoms of NDDs, such as intellectual disability and autism, are first recognized in the toddler stage, circuit differences are likely present at birth and may begin even earlier (Robertson and Baron-Cohen, 2017). From a therapeutic perspective identifying the earliest circuit changes in NDDs is critical because early interventions are more likely to redirect the trajectory of neural development before it is irreversibly changed by genetic and/or environmental factors.

Differences in GABAergic inhibition and excitability have been implicated in the origins of NDDs and autism, and proposed as targets for therapy (Contractor et al., 2021; 2015; Marín, 2016; Rubenstein and Merzenich, 2003). However, the prevalent notion that an imbalance in excitatory and inhibitory signaling is associated with NDDs is principally based on observations in adulthood. In the last decade, as our understanding of cortical development grew significantly, there has been increased awareness about the important developmental role of inhibitory interneurons (INs) in shaping circuits (Dorrn et al., 2010; Fishell and Kepecs, 2020). The proper density, function, and integration of INs into cortical networks all depend on genetic and activity-dependent programs (Wamsley and Fishell, 2017). Deviation from the typical trajectory of these developmental programs in NDDs could have an impact on functional circuit assembly (Contractor et al., 2021; Marín, 2016). For example, hypofunction of cortical INs has been described in multiple models of autism and other psychiatric conditions (Chen et al., 2020; Contractor et al., 2021; Marín, 2012; Salazar et al., 2018), but the nature of GABAergic population dynamics throughout neonatal development in NDDs remain an unexplored territory.

To investigate this, we focused on Fragile X Syndrome (FXS) because it is the most common single gene cause of intellectual disability and autism (Rifé et al., 2003), and because hypoactivity of parvalbumin (PV)-expressing INs has been observed repeatedly in *Fmr1^−/−^* knockout mice, the principal animal model of FXS (Antoine et al., 2019; Berzhanskaya et al., 2017; Domanski et al., 2019; Goel et al., 2018). We show that PV-INs and their precursors, which originate in the medial ganglionic eminence (MGE) and express the transcription factor Nkx2.1, are hypoactive in the primary somatosensory cortex (S1) as early as postnatal day (P) 6 and fail to modulate excitatory neurons in *Fmr1^−/−^* mice. Their density is also reduced from early postnatal development to adulthood in *Fmr1^−/−^*mice and, remarkably, also in humans with FXS. An early intervention with Designer Receptor Exclusively Activated by Designer Drugs (DREADDs) to increase Nkx2.1-IN firing at P5-P9 in *Fmr1^−/−^*mice failed to fully restore circuit dysfunction in S1 despite partially correcting the cortical transcriptome and PV-IN density. However, boosting PV-IN activity with a Kv3.1 channel modulator after the S1 critical period (P15-P20) significantly improved both circuit and behavioral sensory phenotypes of *Fmr1^−/−^* mice. Thus, circuit changes in FXS (and perhaps in other NDDs) can be reversed by targeting PV-INs, but the timing of circuit interventions may be critical.

## RESULTS

### Reduced activity of cortical PV- INs and MGE-derived INs in early postnatal *Fmr1*^−/−^ mice

Several in vivo studies have shown that the activity of cortical PV-INs is reduced in adult *Fmr1*^−/−^ mice (Antoine et al., 2019; Goel et al., 2018). Hence, we previously discovered pronounced circuit changes in S1 during the critical period at P14-P16 Gonçalves et al., 2013; He et al., 2017), that could be due to reduced inhibition, including fewer whisker-responsive excitatory pyramidal (Pyr) neurons and lack of neuronal adaptation. Therefore, we used in vivo two-photon calcium imaging (see Methods) at P15 to record from PV-INs in layer (L) 2/3 of the S1 barrel field of PV-Cre^+/−^ [WT or *Fmr1*^−/−^] mice injected with AAV1-CAG-Flex-GCaMP6s (**Fig 1A-C**). We found that both spontaneous and whisker-evoked activity of PV-INs were significantly reduced by ∼35% in *Fmr1*^−/−^ mice (n=7) compared to WT mice (n=6) (**Fig. 1D-F**). Although the percentage of active PV-INs during spontaneous activity was significantly reduced (**Fig. 1E**), the percentage of whisker-responsive PV-INs was not (**Fig. 1F**). Somatostatin-expressing INs (SST-INs) and PV-INs both arise from a common precursor in the MGE (Marques-Smith et al., 2016; Pouchelon et al., 2021; Tuncdemir et al., 2016). Therefore, we also imaged the activity of SST-INs in vivo at P15 in *Sst*- FlpO^+/−^ mice injected with AAV8-Ef1a-fDIO-GCaMP6s (**Fig.1SA,B**). We did not find significant differences between genotypes in either spontaneous or whisker-evoked activity (**Fig.S1C-E**). Thus, PV-INs in S1, but not SST-INs, manifest significant hypoactivity in two-week-old *Fmr1*^−/−^ mice.

**Figure 1:**
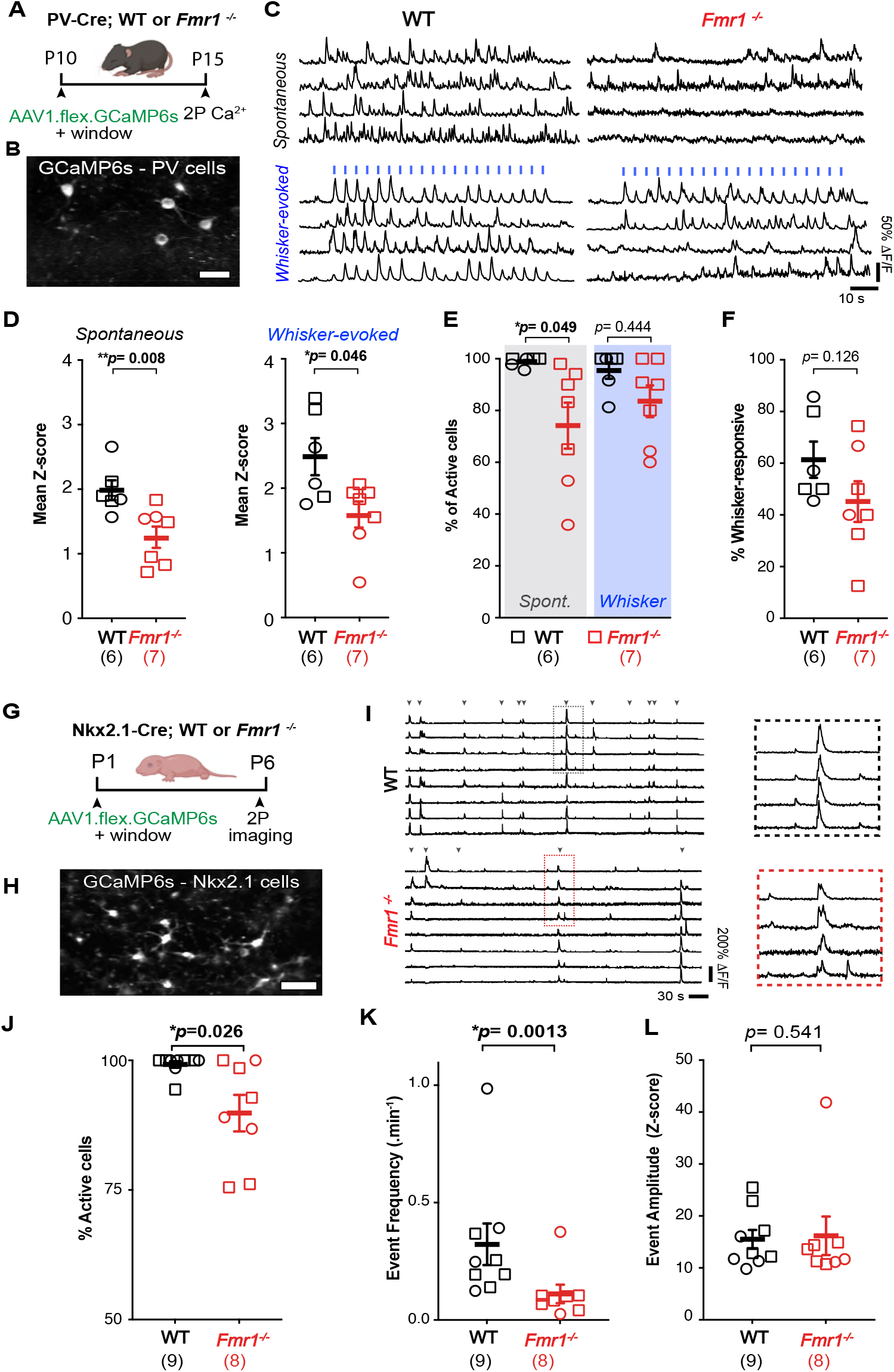
PV-INs and their MGE-derived precursors are hypoactive in S1 of developing *Fmr1^−/−^* mice. a. Cartoon of experimental design for calcium imaging recordings in PV-INs. b. Example field of view of PV-INs expressing AAV1-flex-GCaMP6s in S1 cortex of PV-Cre mice (scale bar= 25μm). c. Representative traces of PV-IN calcium transients during spontaneous and whisker-evoked recordings in WT and *Fmr1*^−/−^ mice. Vertical blue bars represent epochs of whisker stimulation (1 s at 10 Hz, 3 s i.s.i.). d. Mean Z-scores of spontaneous (left) and whisker-evoked activity (right) of PV-INs at P15 are significantly lower in *Fmr1*^−/−^ than in WT mice. In panels d-g and j-k, symbols represent individual mice (sample size in parenthesis), with females and males represented by circles and squares, respectively. (spontaneous: 1.98 ± 0.15 for WT vs. 1.28 ± 0.17 for *Fmr1*^−/−^; whisker-evoked: 2.49 ± 0.29 for WT vs. 1.61 ± 0.21 for *Fmr1*^−/−^, respectively; p= 0.008 and p= 0.046, M-W t-test, n=6 WT and n=7 *Fmr1*^−/−^ mice). e. Percentage of active PV-INs in WT and *Fmr1*^−/−^ mice (spontaneous: 98.9 ± 0.8% for WT vs. 74.2 ± 8.9% for *Fmr1*^−/−^; p= 0.049, whisker-evoked: 95.5 ± 3.2% for WT vs. 83.6 ± 6.1% for *Fmr1*^−/−^, p= 0.444, M-W t-test). f. Percentage of PV-INs that respond to repetitive stimulation in a stimulus-locked manner (see Methods). (61.4 ± 6.9% for WT vs. 45.2 ± 7.9% for *Fmr1*^−/−^; p= 0.126, M-W t-test). g. Experimental design for in vivo recordings at P6 in *Nkx2.1*-Cre mice. h. Example field of view of Nkx2.1-INs expressing AAV1-flex-GCaMP6s in S1 cortex *Nkx2.1*-Cre mice (scale bar= 25μm). i. Example calcium traces of spontaneous activity of Nkx2.1-INs in WT and *Fmr1*^−/−^ mice. Inset shows expanded traces for representative synchronous network events. j. The percentage of active Nkx2.1-INs at P6 was significantly lower in *Fmr1*^−/−^ mice than in WT controls. (99.2 ± 0.6% for WT and 89.8 ± 3.5% for *Fmr1*^−/−^, n= 9 and 8, respectively; p= 0.026, MW t-test). k. The frequency of synchronous network events for Nkx2.1-INs was significantly lower in *Fmr1*^−/−^ mice. (0.32 ± 0.09 events per min for WT vs. 0.11 ± 0.04 for *Fmr1*^−/−^, n=9 WT and n=8 *Fmr1*^−/−^, p= 1.4×10^−3^, MW *t*-test). l. The amplitude of calcium transient events of Nkx2.1-INs was not different between genotypes (15.5 ± 1.8 for WT vs. 16.2 ± 3.7 for *Fmr1*^−/−^, p= 0.541, MW t-test).

During the period that spans between the establishment of barrels and the closure of critical period, prospective GABAergic perisomatic circuits display early structured population dynamics that plays a role in shaping developing barrel cortex (Mòdol et al., 2020; Stern et al., 2001). We next investigated whether early GABAergic interneuron activity during the first neonatal week contributes to the previously described early signs of cortical circuit dysfunction (e.g., excess synchrony in neonatal mice) (La Fata et al., 2014). Cortical PV-INs do not express PVALB before P10 (Butt et al., 2008; de Lecea et al., 1995; Pouchelon et al., 2021), but their precursors from the MGE can be identified through their expression of the transcription factor *Nkx2.1*. We used in vivo calcium imaging to record from MGE-INs at P6 in Nkx2.1-Cre mice injected with AAV1-CAG-Flex-GCaMP6s into S1 at P1 (**Fig. 1G-I)**. Cortical network activity in early postnatal mice is dominated by large and infrequent synchronous network events that propagate as waves across the cortex (Golshani et al., 2009; Rochefort et al., 2009; Yuste et al., 1992). We confirmed that MGE-INs participated in synchronous cortical activity, as previously reported (Duan et al., 2020). However, we found a significantly lower proportion of active Nkx2.1-INs (**Fig. 1J**) and a lower frequency of calcium transients (reduced by ∼65%; **Fig. 1K**) in *Fmr1*^−/−^ mice (n=8) compared to WT controls (n=9), even though the amplitude of events was similar (**Fig. 1L**). Hence, Nkx2.1-INs (precursors of PV and SST-INs) are already hypoactive at P6 and less likely to participate in synchronous network activity.

### Future PV-INs within the Nkx2.1-IN population fail to modulate Pyr cells in neonatal *Fmr1*^−/−^ mice

The hypoactivity of Nkx2.1-INs could account for the previously reported hypersynchrony and hyperactivity of Pyr neurons in early postnatal *Fmr1*^−/−^ mice Gonçalves et al., 2013; La Fata et al., 2014). Indeed, the maturation of cortical network activity depends on the proper integration and function of GABAergic INs (De Marco García et al., 2011; Fishell and Kepecs, 2020; Micheva and Beaulieu, 1995) and co-activation of INs and Pyr cells during the first postnatal week is thought to restrict the spread of spontaneous synchronous cortical network events (Duan et al., 2020; Leighton et al., 2021). However, even though GABAergic perisomatic axons are observed in the first postnatal days, functional synaptic inhibition in S1 does not emerge until P8-10 (Favuzzi et al., 2019; Gour et al., 2021). To understand the functional impact of Nkx2.1-IN hypoactivity on early network activity in *Fmr1*^−/−^ mice, we used an all-optical two-photon optogenetic approach coupled with an intersectional genetic strategy (Fenno et al., 2020) to activate future PV-INs at P10 without activating SST-INs (see Methods). To this end, we generated *Nkx2.1-Cre^+/−^*;*Sst- FlpO^+/−^* mice and injected rAAV8-nEF1-Con/Foff-ChRmine-oScarlet (allowing expression of ChRmine in *Nkx2.1*-Cre^+^/*Sst*-Flp^−^ cells) together with AAV1-CAG-Flex-GCaMP6s and AAV1.Syn.GCaMP6s into S1 at P1 (**Fig. 2A,B**). This all-optical approach allowed us to record from both INs and Pyr cells simultaneously at P10 while we delivered pulses of laser light (1 s pulses at 1,040 nm every 3 s) (**Fig. 2C,D)**. Although laser stimulation significantly drove the activity of *Nkx2.1^+^;Sst ^−^* -INs in both WT and *Fmr1*^−/−^ mice (**Fig. 2C-E**), it only reduced Pyr cell activity and the proportion of active cells in WT mice, as manifested by a lower frequency and amplitude of synchronous events (**Fig. 2F, Fig. S2A,B**). Laser stimulation also significantly decorrelated activity in both WT mice and *Fmr1*^−/−^ mice, but the magnitude of the effect was negligible in the latter (**Fig. 2G, Fig. S2C**). As expected, there was no effect on network dynamics in control mice that did not express ChRmine (**Fig. S2D,E**). Taken together, these results show that MGE-INs, specifically the precursors of PV-INs, are hypoactive in *Fmr1*^−/−^ mice at P6-P10 and, importantly, they fail to properly modulate Pyr network activity.

**Figure 2:**
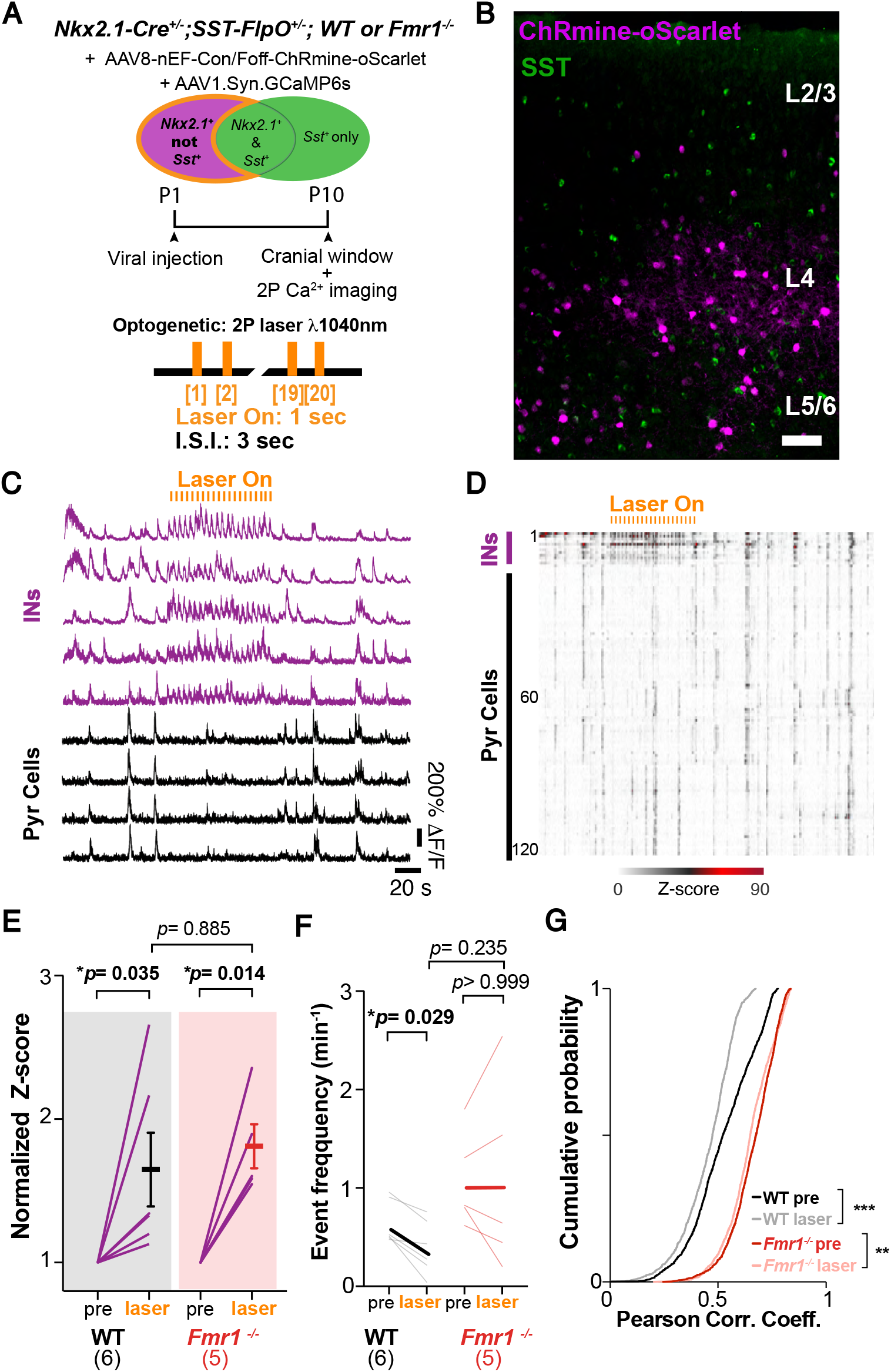
Nkx2.1-Cre^+^ INs form a weak functional network with Pyr cells in neonatal *Fmr1*^−/−^ mice. a. Experimental design for optogenetic experiments. *Nkx2.1*-Cre mice (*Fmr1*^−/−^ or WT) were injected with a CreOn/FlpOff-Chrmine virus at P1 to express the opsin Chrmine in Nkx2.1-INs but not in SST-INs. Calcium imaging was done at P10 before, during, and after 20 orange light pulses (1 s- long, 3 s i.s.i.; 1,040 nm). b. Coronal section through S1 from *Nkx2.1-Cre^+/−^;Sst-FlpO^+/−^*mouse at P10-11 immunostained for SST (green) and showing expression of ChRmine-oScarlet (magenta) showing no overlap between them. c. Representative calcium traces for 5 presumed *Nkx2.1^+^/Sst^−^* INs (magenta) and 4 Pyr cells (black). d. Raster plot of neuronal activity in a representative WT mouse. Presumed *Nkx2.1*^+^/*Sst*^−^-INs are grouped at the top. Note how the frequency (and amplitude) of synchronous network events is reduced for a substantial proportion of Pyr cells during optogenetic stimulation (Laser On). e. Mean z-score of activity in *Nkx2.1^+^/SST^−^* INs before (pre) and during optogenetic stimulation (laser) in WT and *Fmr1*^−/−^ mice. Each line in panels E-F represents an individual mouse (mean normalized z-score increased by 63.7% ± 25.5 for WT, p= 0.035, and 79.9% ± 15.3, p=0.014, for *Fmr1*^−/−^ mice upon laser-on stimulation; two-way ANOVA post-hoc Tukey test; n=6 and 5, respectively). f. Mean frequency of Pyr cell calcium transients was significantly lower during optogenetic stimulation in WT mice but was unchanged in *Fmr1*^−/−^ mice. (0.63 ± 0.09 events.min^−1^ pre vs. 0.38 ± 0.11 with laser; p= 0.029 in WT; 1.06 ± 0.21 events.min^−1^ pre vs. 1.06 ± 0.43 with laser for *Fmr1*^−/−^, p> 0.99; two-way ANOVA, post-hoc Tukey). g. Pair-wise correlation coefficients of Pyr cells were significantly modulated by optogenetic stimulation of *Nkx2.1*^+^/*Sst*^−^ INs in both WT and *Fmr1*^−/−^ mice but the magnitude of the effect was greater in WT (mean corr coeff WT: 0.52 ± 0.00 for pre vs. 0.45 ± 0.00 for laser; p= 2.2 x 10 ^−16^; *Fmr1*^−/−^: 0.66 ± 0.00 for pre vs. 0.64 ± 0.00 for laser, p= 3.3 x 10 ^−9^; Kolmogorov-Smirnov test).

What are the consequences of having Nkx2.1-INs that are hypoactive and largely decoupled from Pyr cells during neonatal cortical development? It is known that a wave of programmed cell death affects MGE-INs from P6 to P10 in mice (Southwell et al., 2012), which is regulated by cortical network activity(Duan et al., 2020; Wong et al., 2018). Moreover, decreasing or increasing the activity of INs in the neonatal period, results in lower or higher density of INs in juvenile mice, respectively (Denaxa et al., 2018; Priya et al., 2018). We considered the possibility that the significant hypoactivity of Nkx2.1**-**INs in *Fmr1^−/−^* mice leads to their excessive cell death and, eventually, a reduced density of PV-INs. Previous reports of reduced PV-IN density in adult S1 and in juvenile primary auditory cortex of *Fmr1*^−/−^ mice had used PVALB immunoreactivity (Selby et al., 2007; Wen et al., 2018). However, because PVALB expression correlates with PV-IN activity levels (Donato et al., 2013), we chose instead to quantify their density in *PV*-Cre;tdTom mice (see Methods). We found a drastic reduction in the density of tdTom^+^ PV-INs in *Fmr1*^−/−^ mice at P15 throughout the cortex (**Fig. S3A**), including a 65% reduction in S1 across both infra- and supragranular layers (**Fig. 3A,B**). A significant reduction in PV-IN density was also observed in S1 of 9-10 month-old *Fmr1*^−/−^ mice (**Fig. S3B**), suggesting their loss is permanent.

**Figure 3:**
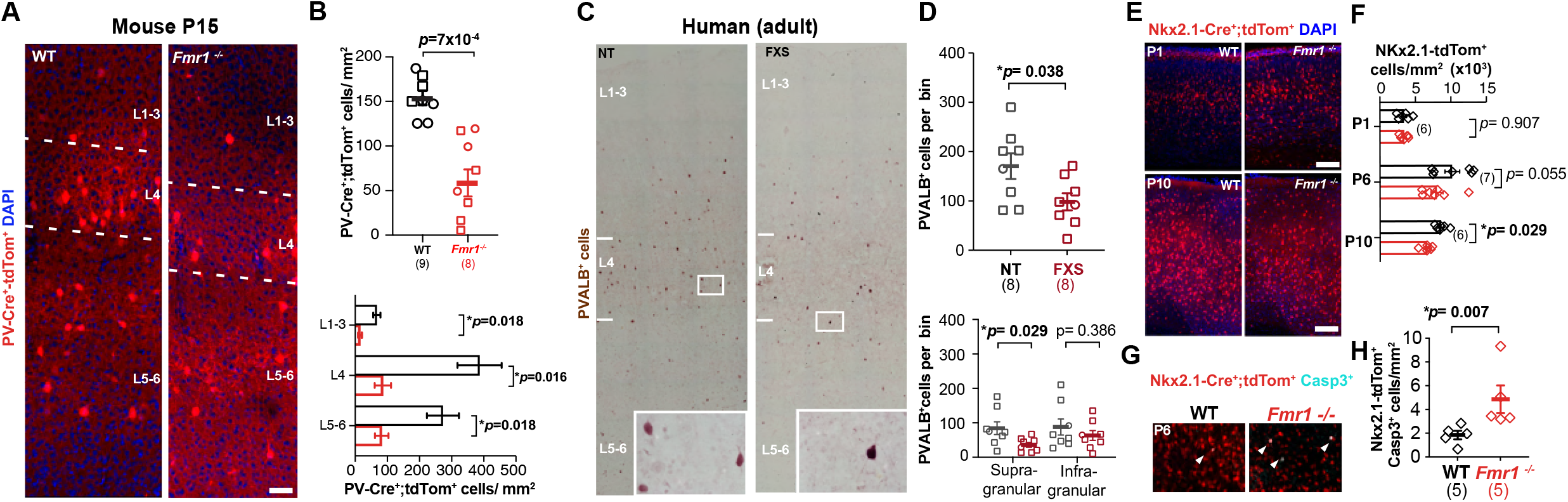
Reduced density of PV-INs and their MGE-derived precursors in mice and humans with FXS. **a.** Example images from coronal sections through barrel field of S1 from *PV*-Cre;tdTom mice (WT and *Fmr1*^−/−^) at P15. Blue denotes nuclear staining with DAPI. Scale bar: 50 μm. **b.** Mean density of *PV*-tdTom*+* INs is significantly lower in *Fmr1^−/−^* mice overall (top) and across cortical layers (bottom). Symbols in panels B, D, and F represent individual mice (sample size in parenthesis), with females and males represented by circles and squares, respectively (diamonds in F indicate the sex was not recorded). (total density: 154.1 ± 8.0 cells per mm^2^ for WT vs. 58.5 ± 15.1 for *Fmr1*^−/−^; n=9 and n= 8, respectively; p= 0.0007, unpaired *t*-test; L2/3: 56.1 ± 9.4 vs 12.7 ± 4.5, p= 0.018; L4: 322.0 ± 57.4 vs. 71.7 ± 21.1, p= 0.016; L5/6: 227.3 ± 41.5 vs. 68.7, p= 0.018 two-way ANOVA post-hoc Holm-Sidak test,). **c.** Example images from coronal sections through BA3 (somatosensory cortex) from neurotypical (NT) and FXS human tissue showing reduced PVALB immunoreactivity in FXS cases across cortical layers. Scale bar: 100 μm. **d.** Mean density of PVALB+ cells in BA3 is significantly lower in FXS cases overall (top) and in supragranular cortical layers (bottom). (170.6 ± 26.0 PVALB+ cells per bin for NT vs. 98.1 ± 17.4 for FXS cases, n=8 cases each; p=0.038, unpaired *t*-test) (supragranular layer: 83.6 ± 18.6 for neurotypical vs 35.5 ± 7.0, p=0.0294; infragranular: 87.0 ± 23.2 vs. 62.6 ± 14.3, p=0.386, unpaired *t*-tests). **e.** Coronal sections through S1 showing *Nkx2.1*-Cre;tdTom^+^ INs (red) in WT and *Fmr1*^−/−^ mice at P1 and P10 (DAPI: blue). Scale bar: 100 μm. **f.** Mean density of Nkx2.1;tdTom+ INs is lower in *Fmr1*^−/−^ mice at P6 and P10, but not at P1. (P1: 3,408 ± 354 cells/mm^2^ for WT vs. 3,472 ± 548 for *Fmr1*^−/−^, p=0.907; P6: 10,200 ± 1,027 cells/mm^2^ for WT vs. 8,035 ± 860 for *Fmr1*^−/−^, p= 0.056; P10: 8,633 ± 318 for WT vs. 6,650 ± 279 for *Fmr1*^−/−^, p=0.029; n=6 per genotype at P1 and P10, n=7 per genotype at P6, two-way ANOVA with post-hoc Holm-Sidak). **g.** Coronal sections through S1 from *Nkx2.1-Cre* mice (WT and *Fmr1*^−/−^) at P6 immunostained for cleaved Caspase-3 (cyan) showing rare double-labeled cells (arrowheads). Scale bar: 100 μm. **h.** Mean density of *Nkx2.1*-tdTom^+^ INs that express Caspase-3 was significantly higher in *Fmr1*^−/−^ mice. (1.85 ± 0.35 cells/mm^2^ for WT vs. 4.87 ± 1.16 for *Fmr1*^−/−^, p=0.007, M-W test).

To determine whether this phenotype is perhaps unique to mice, we also quantified the density of three major types of INs in human autopsy material from adult FXS cases and age-matched controls (n=8 each) using immunohistochemistry (see Methods). Remarkably, we found a significant 42% reduction in PVALB-expressing INs in Brodmann area 3 (equivalent to S1 in mice) in FXS cases (**Fig. 3C,D**). In contrast, the density of calretinin-expressing and calbindin-expressing IN subclasses was not different between FXS cases and controls (**Fig. S4**).

We next examined the density of *Nkx2.1Cre*^+^;tdTom^+^ INs throughout early postnatal development in mouse S1 (**Fig. 3E**). A reduced density of Nkx2.1-INs was seen in *Fmr1*^−/−^ mice at P6 and P10 compared to WT controls (**Fig. 3F**), but not at P1, suggesting that the lower density of PV-INs is not caused by reduced migration and/or neurogenesis of MGE-INs. To confirm that the lower density of MGE-INs reflects increased levels of developmental apoptosis, we immunostained brain sections for the apoptosis marker Caspase-3 (Southwell et al., 2012). We found a significantly higher density of Caspase-3 expressing MGE-INs at P6 in *Fmr1*^−/−^ mice compared to WT controls (n=5 each; **Fig. 3G,H**). Hence, the low density of PV-INs in humans with FXS and in *Fmr1*^−/−^ mice is evident at the earliest stages of cortical development, several days before they assume their PVALB+, fast-spiking identity.

The absence of the Fragile X Protein in FXS triggers profound transcriptional dysregulation (Darnell and Klann, 2013; Miyashiro et al., 2003; Sharma et al., 2019). Because the hypoactivity of Nkx2.1-INs in *Fmr1*^−/−^ mice is one of the earliest developmental brain phenotypes, it likely triggers significant gene dysregulation in FXS (Dehorter et al., 2015; Mahadevan et al., 2021; Wamsley and Fishell, 2017). We performed mRNA sequencing of S1 cortical samples at P15 to uncover putative molecular changes associated with PV-IN dysfunction in *Fmr1*^−/−^ mice and to determine whether artificially activating *Nkx2.1^+^*-INs via DREADDs during neonatal development might restore gene expression to WT levels. We used a Ribotag approach (Sanz et al., 2009) that allowed us to assess transcriptional changes not only in Nkx2-1-INs but also globally in S1 (Mahadevan et al., 2020) (see Methods; **Fig. 4A,B**). First, we generated *Nkx2.1-Cre^+/−^*;*Rpl22^HA/-^* [WT or *Fmr1^−/−^*] triple transgenic mice and virally expressed hM3Dq (or mCherry as a control) in Nkx2.1-INs from P1 onward. Following chronic administration of the DREADD agonist compound 21 (Thompson et al., 2018) (C21, 1 mg/kg, s.c., twice daily) from P5 to P9, we performed RNA sequencing at P15 (**Fig. 4A**). We identified 2,898 differentially expressed (DE) genes in the bulk RNA samples of *Fmr1*^−/−^-mCherry mice (n=6 mice), as compared to WT-mCherry mice (n=8 mice) (**Fig. 4C,E**). Gene ontology analysis (see Methods) revealed an enrichment in genes involved in “*Synapse organization*” and “*Neuron* a*poptotic process*” among other categories (**Fig. 4C**). Notably, we found that several markers for MGE-INs (Favuzzi et al., 2019; Mahadevan et al., 2021; Paul et al., 2017) were significantly downregulated in *Fmr1*^−/−^ mice, including several PV-specific genes (e.g., *Pvalb, Bcan, Syt2, Kcnab3*), while other MGE-IN genes were upregulated, including some SST-specific genes (*Sst*, *Pcdh18*) (**Fig. 4C,H**). These patterns of gene expression (downregulation of PV-IN genes, upregulation of neuronal apoptosis genes) are consistent with the reduced activity and density of PV-IN we observed at P15 in *Fmr1*^−/−^ mice (**Figs. 1-3**).

**Figure 4:**
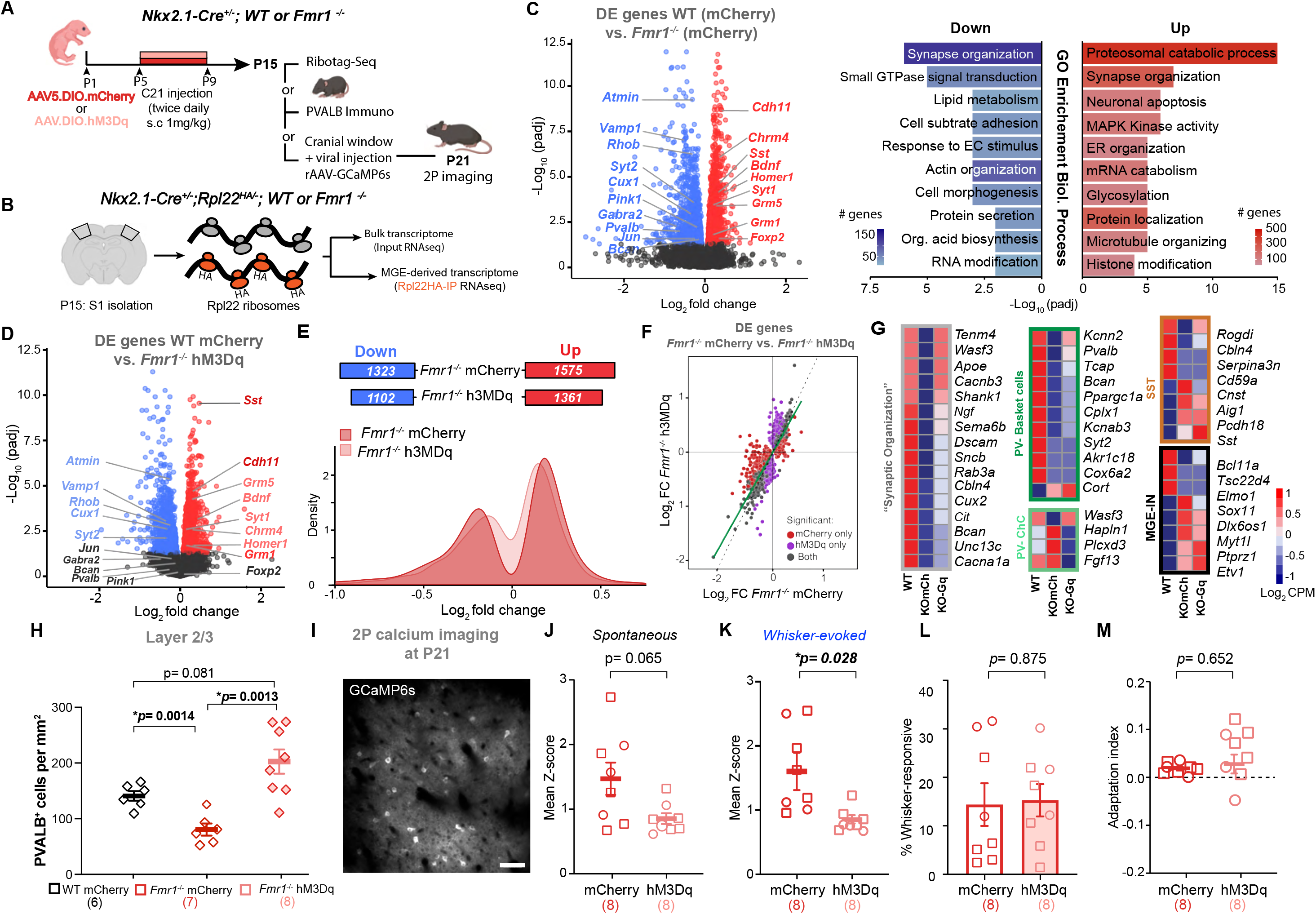
Chronic chemogenetic activation of Nkx2.1-INs mitigates PV-IN loss and gene dysregulation in neonatal *Fmr1*^−/−^ mice, but not S1 circuit dynamics. a. Experimental design for chronic chemogenetic activation of Nkx2.1-INs (from P5 to P9) in *Fmr1*^−/−^ mice to assess Nkx2.1-specific Ribotag RNA-seq at P15, density of PVALB+ INs at P15, and in vivo 2P calcium imaging at P21. b. Cartoon of Ribotag approach for RNA-seq in both bulk samples and pull-down. c. Left: Volcano plot of differentially expressed (DE) genes in the bulk RNA of *Fmr1*^−/−^-mCherry mice (n=6) compared to WT-mCherry control group (n=8). A few genes are annotated (because of their role in IN and synapse function) amongst significantly downregulated (blue) or upregulated genes (red). Right: Top 10 ‘GO terms’ using the biological process package (see Methods). d. Volcano plot of DE genes in the bulk RNA of *Fmr1*^−/−^-hM3Dq mice (n=7) compared to WT-mCherry controls (n=8). Genes from c that were fully or partially corrected by the DREADD manipulation are in gray or light color font, respectively. e. Top: The total number of down- and up-regulated genes (compared to WT-mCherry controls) was smaller in *Fmr1*^−/−^-hM3Dq mice than in *Fmr1*^−/−^-mCherry. The expression of 435 genes was ‘corrected’ by the chronic chemogenetic treatment in *Fmr1*^−/−^-hM3Dq mice. Bottom: Density plot showing how the log_2_ fold change was affected by the DREADD manipulation (fewer genes are different from WT mice in the *Fmr1*^−/−^-hM3Dq group). f. Correlation plot of genes affected in the *Fmr1*^−/−^-mCherry and *Fmr1*^−/−^-hM3Dq. Red and purple dots denote DE genes that are uniquely and significantly different from WT controls in *Fmr1*^−/−^-mCherry and *Fmr1*^−/−^-hM3Dq mice, respectively. Grey dots are shared DE genes in both groups. Note that purple genes are on average less different from WT than red genes. g. Representative examples of how DREADD manipulation affected genes for the “synapse organization” GO term (gray), or different subclasses of MGE-INs (dark green: basket cells; light green: Chandelier cells; brown: SST cells; black: global MGE-IN markers. Scale bar represents the log_2_ CPM (count per millions). h. Quantification of total PVALB^+^ cell density at P15 in the barrel field of S1 of WT-mCherry, *Fmr1*^−/−^- mCherry and *Fmr1*^−/−^-h3MDq mice. (160.9 ± 6.5 cells/mm^2^, 87.4 ± 5.5, and 118.6 ± 9.0, n=6, 7, and 8, respectively). i. Example field of view of Pyr neurons expressing AAV-GCaMP6s in S1 of *Nkx2.1-Cre^+/−^*;*Fmr1*^−/−^ (scale bar= 100 μm). j. Mean Z-scores for spontaneous activity of Pyr cells are lower in *Fmr1*^−/−^-hM3Dq mice compared to *Fmr1*^−/−^-mCherry mice after chronic C21 injections. (spontaneous: 1.47 ± 0.25 vs. 0.85 ± 0.08, p=0.065, n=8 per group) k. Mean Z-scores for whisker-evoked activity (right) of Pyr cells are significantly lower in *Fmr1*^−/−^- hM3Dq mice compared to *Fmr1*^−/−^-mCherry mice after chronic C21 injections. (1.58 ± 0.29 vs. 0.84 ± 0.06, p=0.028, MW t-test, n=8 per group). l. The proportion of whisker-responsive Pyr cells was not changed by DREADDs in *Fmr1*^−/−^ mice. (14.4 ± 4.4% vs. 15.3 ± 3.3%, p=0.875, unpaired t-test) m. The neuronal adaptation index of Pyr cells to repetitive whisker stimulation was not changed by chemogenetics. (0.19 ± 0.00 for WT vs. 0.03 ± 0.01 for *Fmr1*^−/−^; p=0.652, unpaired t-test).

We next compared the bulk transcriptome of *Fmr1*^−/−^-hM3Dq mice (n=7 mice) treated with C21 from P5-P9 to that of similarly treated *Fmr1*^−/−^-mCherry controls and found 15% fewer DE genes (2,463 vs. 2,898; **Fig. 4D-F, Fig. S5**). Interestingly, the expression of several genes enriched in MGE-INs was restored to WT levels, including genes specific to basket and chandelier cells (*Pvalb, Kcnn2, Wasf3, Sox11*), as well as SST-INs (*Cbln4, Cnst)* **(Fig. 4G)** (Favuzzi et al., 2019; Mahadevan et al., 2021; Paul et al., 2017). Genes whose expression was rescued by Gq-DREADDs belong primarily to *Synapse organization*, *RNA metabolism*, and *Neuron* a*poptotic process*, among others (**Fig. 4G, Figs. S5 & S6**). On the other hand, other sets of genes either remained unchanged or were newly dysregulated by Gq-DREADDs (**Fig. 4G, Fig. S6**).

As far as the Nkx2.1-specific transcriptome (RiboTag ‘pull-down’), we identified 2,039 DE genes samples from *Fmr1*^−/−^-mCherry mice, compared to WT-mCherry mice, including once again *Pvalb*, *Pink1* and *Gabra2* among the down-regulated group (**data Fig. S7**). GO enrichment analysis revealed that the most up-regulated genes belong to “*Synapse organization*” and “*Cognition*” categories, but we did not identify GO terms for down-regulated genes (**Fig. S7B**). Unexpectedly, we identified a greater number of Nkx2.1-specific DE genes (6,105) in *Fmr1*^−/−^- hM3Dq mice treated with C21, compared to WT controls, and the number of genes in the “*Synapse organization*” category was higher than in *Fmr1*^−/−^-mCherry mice (**Fig. S7C-E**). hM3Dq + C21 treatment had only a modest effect in genes associated with MGE-INs, and generally caused further dysregulation compared to WT mice (**Fig. S7F).**

In summary, chronically increasing the activity of Nkx2.1-INs in *Fmr1*^−/−^ mice from P5 to P9 (during their period of developmental apoptosis and synaptogenesis with Pyr cells) can partially correct the cortical transcriptome, although not the MGE-specific transcriptome.

### Activating of *Nkx2.1^+^*-INs in neonatal *Fmr1*^−/−^ mice partially restores PV cell density, but not S1 circuit dysfunction

Our findings of reduced firing and density of INs in *Fmr1*^−/−^ mice (**Figs. 1-3**) are consistent with a model in which hypoactivity of the Nkx2.1-IN population (and their uncoupling to Pyr cells) results in excess programmed cell death of future PV-INs. We reasoned that artificially increasing the activity of Nkx2.1-INs in neonatal mice, during the apoptosis window (P5-P10), might rescue the density of PV-INs and perhaps also other circuit phenotypes of older *Fmr1*^−/−^ mice. To test this, we injected rAAV5-DIO-hM3Dq-mCherry (or an rAAV5-DIO-mCherry control virus) in Nkx2.1- Cre;*Fmr1*^−/−^ mice and administered C21 (1 mg/kg, s.c., twice daily) from P5 to P9 (**Fig. 4A**). We found that the density of PVALB-expressing INs in L2/3 at P15 was fully restored to WT levels in hM3Dq mice than in mCherry controls (**Fig. 4H)**, although the effect was less pronounced in deeper layers (**Fig. S8B**). The fact that increasing MGE-IN activity can prevent their death in *Fmr1^−/−^* mice further supports the notion that many MGE-INs eventually die as a result of their hypoactivity.

We next tested whether the same DREADD manipulation in Nkx2.1-INs could restore circuit changes of *Fmr1*^−/−^ mice. We focused on two robust S1 phenotypes we previously reported in S1 of *Fmr1*^−/−^ mice at P15, namely the low percentage of whisker-responsive Pyr cells in L2/3 and their lack of adaptation to repetitive whisker stimulation (C. X. He et al., 2017). Once again, we virally expressed hM3Dq (or mCherry alone) in *Fmr1*^−/−^ mice and injected them with C21 from P5 to P9, implanted cranial windows (and injected rAAV1-Syn-GCaMP6s) at P15 and performed in vivo calcium imaging of Pyr cells at P21 (**Fig. 4I**). Even though C21 administration had ceased days earlier, this early DREADD activation of Nkx2.1-INs caused a significant reduction of whisker-evoked activity of Pyr cells (**Fig. 4J**). However, it did not increase the proportion of whisker-responsive neurons, nor their adaptation to 20 sequential bouts of whisker stimulation (see Methods) (**Fig. 4K,L)**. Altogether these DREADD experiments demonstrate that raising the activity of MGE-derived INs in neonatal *Fmr1*^−/−^ mice can partially restore the density of PV-INs and the expression of ∼15% of cortical genes (including important genes involved in IN maturation and function), but fails to improve circuit dynamics in S1.

### Boosting PV-IN activity after the critical period rescues S1 circuit deficits and tactile defensiveness in juvenile *Fmr1*^−/−^ mice

Our neonatal DREADD manipulation likely failed to restore excitatory neuron responsivity and adaptation because MGE-INs are not yet functionally connected with Pyr cells in *Fmr1*^−/−^ mice before P10, as suggested by our optogenetic experiments (**Fig. 2**). Several studies suggest that some aspects of cortical maturation are transiently delayed in *Fmr1*^−/−^ mice, but eventually catch up to WT levels (“Circuit level defects in the developing neocortex of Fragile X mice.,” 2013; Cruz-Martín et al., 2010; Harlow et al., 2010; Q. He et al., 2014), including the morphology and intrinsic properties of fast-spiking IN properties (Nomura et al., 2017). Thus, we next explored whether increasing PV-IN activity after the S1 critical period (when presumably INs are able to modulate Pyr cells) could rescue circuit changes and tactile defensiveness in *Fmr1*^−/−^ mice (C. X. He et al., 2017). To this end, we first used an acute chemogenetic approach to transiently increase PV-IN activity at P15 by expressing hM3Dq (or mCherry) and GCaMP6s in PV-Cre;*Fmr1*^−/−^ mice and recording L2/3 Pyr neurons in vivo at P15, before and 30-40 min after administering a single dose of C21 (1 mg/kg, s.c.) (**Fig. S9A**). Compared to vehicle injection, acute C21 injection resulted in a significant reduction of spontaneous and whisker-evoked activity of Pyr cells in *Fmr1*^−/−^ mice expressing hM3Dq, but not in the mCherry controls (**Fig. S9B,C**). C21 injection also resulted in a significant increase in the percentage of whisker-responsive Pyr cells in the hM3Dq group (**Fig. S9D**). However, there was no effect on neuronal adaptation after a single C21 injection (**Fig. S9E**). Therefore, acutely boosting PV-IN firing with DREADDs at P15 in *Fmr1*^−/−^ mice can rescue some but not all circuit changes associated with tactile defensiveness.

It is possible that our DREADD strategy fell short of full circuit rescue because we only increased PV-IN activity acutely (with a single dose of C21) and because the Gq virus infected only a small fraction of all PV-INs in S1 (**Fig. S8A**). Therefore, we next tested a chronic pharmacologic approach to achieve a more global and longer-lasting increase of PV cell activity in *Fmr1*^−/−^ mice after the critical period. After ∼P10, cortical PV-INs assume their fast-spiking characteristics, in large part due to their expression of Kv3.1 channels. This subclass of voltage-gated potassium channels, responsible for rapid repolarization that enables their fast-spiking behavior, is almost exclusively expressed in PV-INs (Du et al., 1996; Rudy and McBain, 2001). We reasoned that targeting Kv3.1 channels pharmacologically could be used to modulate the firing of PV-INs as a potential treatment for various NDDs. From an identified series of Kv3.1 positive allosteric modulators, we pursued compound AG00563 (1-(4-methylbenzene-1-sulfonyl)- N-[(1,3-oxazol-2-yl)methyl]-1H-pyrrole-3-carboxamide); **Fig. 5A**) because of its favorable target engagement and selectivity profile when compared to other ion channels (unpublished observations). Patch-clamp recordings of identified PV-INs in acute slices from P14-P16 *PV-*Cre^+^- tdTom^+^*;Fmr1^−/−^* mice (see Methods), showed that bath application of AG00563 (10 µM) significantly increased their excitability (**Fig. 5B,C**). These results were independently confirmed at Lundbeck in fast-spiking neurons in acute rat slices. Of note, bath application of AG000563 did not affect the membrane potential or excitability of Pyr cells, nor the input resistance of PV-IN or Pyr cells (**Fig. S10).**

**Figure 5:**
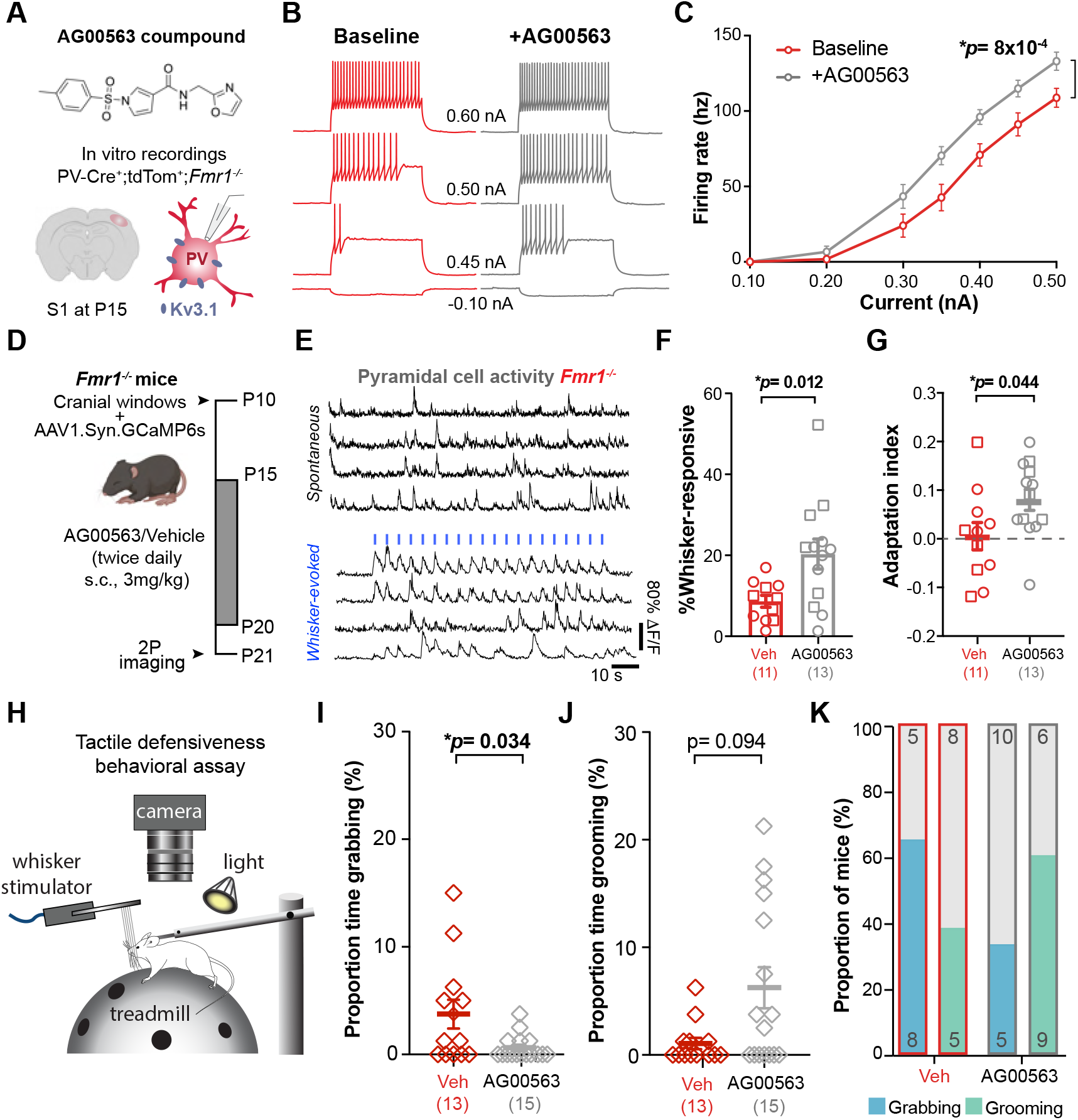
A novel Kv3.1b allosteric modulator, AG00563, ameliorates circuit function and tactile defensiveness in juvenile *Fmr1*^−/−^ mice. a. Chemical structure of the AG00563 compound and experimental design for the in vitro patch-clamp recordings of PV-INs. b. Example traces of action potential trains evoked by 250 ms current injection (50-100 pA steps) in a *PV*-tdTom^+^ cell from a *Fmr1*^−/−^ mouse at P15. Note the increased firing during bath application of AG00563 (gray; 10 µM for 5 min) compared to baseline (red). c. Cumulative input-output curves during baseline (red) or AG00563 (gray) (n= 15 cells from 6 *Pv*- Cre^+/−^;tdTom+;*Fmr1*^−/−^mice, two-way RM ANOVA). d. Experimental design for chronic AG00563 vs. vehicle treatment (5 d from P15 to P20, 3 mg/kg, s.c, twice daily) and in vivo calcium imaging at P21. e. Example traces of L2/3 Pyr cell calcium transients during spontaneous and whisker-evoked recordings in *Fmr1*^−/−^ mouse. Vertical blue bars represent epochs of whisker stimulation (1 s at 10 Hz, 3 s i.s.i.) f. The percentage of whisker-responsive Pyr cells was significantly higher in *Fmr1*^−/−^ mice treated with AG00563 than in vehicle controls. (20.3 ± 3.7% vs. 8.7 ± 1.5%, p=0.012; unpaired *t*-test, n=10 and 13 mice, respectively). g. The adaptation index of Pyr cells was also significantly higher in *Fmr1*^−/−^ mice treated with AG00563-compared to vehicle controls. (0.01 ± 0.03 vs. 0.08 ± 0.03; p=0.0438; unpaired *t*-test). h. Cartoon of tactile defensiveness behavioral assay. i. Proportion of time spent grabbing the stimulator (defensive behavior) was significantly lower in AG00563-treated mice than in vehicle controls (0.62% ± 0.28 s vs. 3.75% ± 1.33 s, p=0.0343, MW t-test, n=15 and 13 *Fmr1*^−/−^ mice, respectively) j. Proportion of time spent grooming was higher in AG00563-treated group, but the difference was not significant (6.25% ± 1.9 s vs. 1.06% ± 0.53 s, p=0.094, MW t-test). k. A smaller percentage of mice showed defensive behavior (grabbing) at least once during whisker stimulation in the AG00563-treated group than among vehicle controls (5/15 mice vs. 8/13, respectively). The opposite was true for adaptive healthy behavior (grooming) (9/15 mice vs. 5/13, respectively).

Using in vivo calcium imaging, we found that acute administration of AG00563 (3 mg/kg, s.c.) at P15 significantly increased the fraction of whisker-responsive Pyr cells, but did not change their adaptation index (**Fig. S9F-H**), which matches results we obtained with acute excitatory DREADDs in PV-INs (**Fig. S9D,E**). We next administered AG00563 (or vehicle) chronically (3 mg/kg, s.c., twice daily) to *Fmr1^−/−^* mice from P15 to P20, and then performed calcium imaging the following day, at least 16 h after the last dose (**Fig. 5D,E**). Thus, by the time we recorded network activity or tested mice behaviorally for tactile defensiveness, any acute effect of the compound was no longer a factor. We found that compared to vehicle controls (n=10 mice) the proportion of whisker-responsive Pyr cells in S1 was significantly higher in AG00563-treated mice (n=13 mice) (**Fig. 5F**), reaching near WT levels (C. X. He et al., 2017). Moreover, the adaptation index of Pyr cells was also significantly increased by AG00563 (**Fig. 5G**). The absence of neuronal adaptation in certain brain circuits is one potential reason why children with NDDs/autism exhibit sensory hypersensitivity, because they are unable to ‘tune out’ non-threatening or non-salient stimuli (Green et al., 2015). We used a tactile defensiveness assay based on repetitive whisker stimulation (C. X. He et al., 2017) (see Methods; **Fig. 5H**) to test whether AG006563 might lessen the maladaptive avoidance/defensive behaviors previously observed in *Fmr1^−/−^* mice. We found that mice chronically treated with AG00563 from P15 to P20 manifested significantly less grabbing during whisker stimulation than vehicle-treated *Fmr1^−/−^* controls (**Fig. 5I**). Moreover, AG00563-treated *Fmr1^−/−^* mice spent more time demonstrating healthy adaptive behaviors, such as grooming (**Fig. 5J**). Not all mice exhibited grabbing or grooming, but overall, more AG00563-treated *Fmr1^−/−^*mice displayed grooming and fewer showed grabbing, at some point during whisker stimulation, while the opposite was true in vehicle-treated *Fmr1^−/−^*mice (**Fig. 5K**). Therefore, chronic pharmacological activation of PV-INs after P15, can ameliorate S1 sensory circuit deficits and rescue behavioral manifestations of tactile defensiveness in juvenile *Fmr1^−/−^* mice.

## DISCUSSION

We set out to identify when IN hypofunction first begins in *Fmr1^−/−^*mice to better understand how developmental trajectories of cortical circuits are changed in FXS and in other NDDs that share a deficit of cortical INs. We used an intersectional strategy to express chemo- and optogenetic tools to manipulate Nkx2.1-INs, as well as in vivo calcium imaging to record from them and their excitatory Pyr cell counterparts. We discovered that: 1. PV-INs and their precursors from the MGE in *Fmr1^−/−^*mice are hypoactive as early as P6 and their density is reduced in both mice and humans with FXS, due to an excess of developmental apoptosis; 2. Profound alterations of the transcriptomic landscape pertaining to cortical circuit organization are present at the closure of the S1 critical period in *Fmr1^−/−^* mice; 3. Increasing the activity of MGE-INs from P5 to P9 partially restores the density of PV-INs and a subset of DE cortical genes, but fails to fully rescue circuit dynamics (likely because MGE-INs fail to modulate neonatal Pyr cell network activity at that stage); 4. In contrast, boosting PV-IN activity with a novel Kv3.1 allosteric modulator after the S1 critical period (P15-P20) does correct network activity in S1 and rescues tactile defensiveness in *Fmr1^−/−^* mice.

### Critical developmental role of Nkx2.1-INs in sensory circuits in FXS

GABAergic INs govern crucial steps in the scaffolding and maturation of brain circuits and have been hypothesized to play a key role in NDDs (Contractor et al., 2021; Marín, 2016). Previous studies have shown that, during the early postnatal period, spontaneous Pyr cell activity is hypersynchronous in *Fmr1^−/−^*mice Gonçalves et al., 2013; La Fata et al., 2014). Our present studies (**Fig. 2G, Fig. S2**), indicate this is not only a robust phenotype, but also the earliest sign of network alteration in FXS (Golshani et al., 2009; Rochefort et al., 2009). Additionally, we now demonstrate that Nkx2.1^+^ INs are hypoactive in neonatal *Fmr1^−/−^* mice during a time that coincides with the emergence of perisomatic GABAergic inhibition onto Pyr cells. Considering the emerging knowledge about IN-Pyr connectivity in neonatal cortex (Anastasiades et al., 2016; Marques-Smith et al., 2016) this hypoactivity is likely to have serious consequences for both structural and functional connectivity, and the eventual processing of sensory inputs. Notably, during typical cortical development, this inhibition is likely important for the desynchronization of network activity, which occurs around P12,(Golshani et al., 2009; Rochefort et al., 2009). Indeed, our all-optical two-photon optogenetic approach demonstrated that activation of Nkx2.1+ INs can drive the decorrelation of Pyr cells in WT mice, which is to our knowledge, the first demonstration of a functional role for INs in this major developmental network transition (**Fig. 2G**). The previously reported developmental delay in this desynchronization in *Fmr1^−/−^*mice (Cheyne et al., 2019; Gonçalves et al., 2013) is likely due to the fact that Nkx2.1+ INs are hypoactive and also functionally connected to Pyr cells. We therefore posit that interventions aimed to restore the typical time course of network desynchronization could be beneficial to ameliorate symptoms of FXS. Our new data suggests that the developmental trajectory of cortical INs begins to deviate very early in individuals with FXS, when highly synchronous neural activity dominates in neocortex (this stage in neonatal mice would roughly correspond to the late 3^rd^ trimester of gestation in humans; (Molnár et al., 2020)). Importantly, our results are not consistent with major disruptions in the generation or migration of INs from the MGE.

### Excess developmental death of PV-INs in FXS

There has been some controversy about whether the lower density of PV-INs reported in several models of NDDs/autism represents an artifact(Contractor et al., 2021). The argument is that those studies had relied on PVALB immunohistochemistry to identify PV-INs in tissue but the problem is that PVALB expression levels are known to correlate with PV-IN activity (Donato et al., 2013); therefore low PV-IN density in autism models merely reflects their hypoactivity (Filice et al., 2016). In the present study we circumvent this potential caveat in *Fmr1*^−/−^ mice by counting tdTom^+^ cells in PV-Cre;Ai9 mice, and confirm a marked reduction in PV-IN density (**Fig. 3A,B**). In some mice the reduction in PV-IN density was so striking that only rare, scattered PV-INs could be identified in the neocortex (**Fig. S3A**, example 2). We also found co-expression of Caspase-3 in a subset of tdTom+ INs during neonatal development, providing some evidence that they succumb to programmed cell death. We also report, for the first time, that the density of PV-INs (but not calretinin- or calbindin-expressing INs) is also reduced in human FXS cases (**Fig. 3C,D**). Of course, this apparent absence of a fraction of PV-INs in human tissue could represent their hypoactivity, or perhaps unique vulnerability of these neurons to the post-mortem delay in FXS. Therefore, it is not yet possible to definitively conclude that PV-IN density is reduced in humans with FXS, but the parallels between mouse and human data in this regard remain striking. Interestingly, organoids derived from human induced pluripotent stem cells from patients with FXS had a lower density of GABAergic neurons (Kang et al., 2021), which was felt to be due to reduced neurogenesis. Because we found a normal density of Nkx2.1+ INs at P1 in *Fmr1*^−/−^ mice, we do not believe that the reduced PV IN density is caused by problems with their birth or migration. Additional studies will be needed to confirm these results in a larger number of human cases, and ideally with attempts to correlate age and symptom severity with PV-IN density in various brain regions.

One of the most striking observations is that, even though we can raise the excitability of INs in neonatal animals, this is not sufficient to restore functional connectivity in S1. We suspect this is because MGE-INs are decoupled from their Pyr cell partners during neonatal development, at least initially. This could reflect changes in INs themselves (e.g., a delay in synaptogenesis), or in post-synaptic Pyr cells (e.g., changes in post-synaptic GABA receptor expression). *Fmr1^−/−^*mice show deficits in Pyr cell synaptic inputs onto FS INs (Gibson et al., 2008; Nomura et al., 2017), and genes involved in “synapse organization” were amongst the most dysregulated in our dataset. Our acute DREADD experiments at P15 suggest that cortical PV-INs in *Fmr1^−/−^* mice are eventually able to establish functional connections with Pyr cell partners because they can inhibit them at P15 (see **Fig. S9B,C**), though they remain hypoactive and their numbers are permanently reduced for their lifetime. However, the maturation of morphology, intrinsic properties, and feedforward connectivity of PV-INs are only delayed in *Fmr1^−/−^* mice (Gibson et al., 2008; Nomura et al., 2017). This might explain why we could restore circuit dynamics in S1 and ameliorate sensory avoidance behaviors in *Fmr1^−/−^* mice by boosting PV-IN firing after P15.

### Implications for treatment of FXS

Because symptoms of NDDs are first recognized in toddlers, it is generally understood that they arise because of changes in the brain that occur even earlier, likely in utero. This makes sense for FXS because expression of the Fragile X Protein in the brain starts prenatally, so its absence could change the typical developmental trajectory of brain maturation in the third trimester of gestation or earlier. Therapeutic interventions for intellectual disability or autism that begin at the earliest stages of brain development should therefore be the most effective (Robertson and Baron-Cohen, 2017; Veenstra-VanderWeele and Warren, 2015). Indeed, in recent gene therapy studies for Angelman syndrome, delayed reinstatement of the *Ube3a* gene in adolescent mice was less effective in reversing deficits in the mouse model than a similar intervention earlier in development (Mei et al., 2016; Silva-Santos et al., 2015). Perhaps there is a lesson to be learned in our different manipulations about the timing of potential PV-targeted therapies in NDDs. Although increasing the activity of Nkx2.1+ INs with DREADDs in neonatal mice did not fully rescue *Fmr1^−/−^*circuit phenotypes in S1, increasing PV-IN firing at P15 (with AG00563 or with Gq-DREADDs), even acutely, was able to increase the fraction of whisker responsive Pyr cells. This offers hope in FXS, because it means that interventions to boost IN activity need not start at birth (this may not be in fact desirable), but could be efficacious if started in childhood, adolescence, or even adulthood (Goel et al., 2018).

Atypical sensory perception is present in toddlers with FXS and other NDDs and is believed to contribute to social communication differences, repetitive behaviors and learning disability in adulthood (Robertson and Baron-Cohen, 2017; Rogers et al., 2003). Our encouraging preclinical results with the Kv3.1 activator AG00563 suggest that increasing activity of PV-INs is a plausible strategy to lessen not only tactile defensiveness in FXS, but perhaps also other symptoms in children and adults with NDDs.

## Acknowledgments

The authors thank Anis Contractor, Aaron McGee for feedback on the manuscript. We also thank Fuying Gao and Riki Kawaguchi form the bioinformatic core for functional genomics common research resource and the Technology Center for Genomics and Bioinformatics core at UCLA.

## Author contributions

N.K. and C.P.-C. conceived the project and designed the experiments with help from V.M.-C. for the human studies and D.M. and B.L. for the in vitro recordings. N.K generated the mouse lines, performed the in vivo imaging, chemogenetic, optogenetic and immunohistology experiments. N.K., A.L. and D.T.C. performed the behavioral experiments and analyzed the in vivo imaging and behavioral data. P.J and V.M-C performed and analyzed the human tissue experiments. N.K., C.P.-C., M.G., and A.S. designed the Ribotag-RNAseq experiments. A.S. generated the Ribotag mouse lines, performed the Ribotag-RNAseq experiments and analysis, with help from N.K., who performed the viral injection and C21 agonist treatment. N.K., C.P.-C., B.L. and D.B. designed and B.L performed the in vitro electrophysiology recordings. N.K., C.P-C, A.G.S., C.M., and B.H. designed the AG00563 experiments. N.K., A.S., D.B., and C.P.-C. interpreted the data. N.K. and C.P.-C. wrote the manuscript with input from other authors.

## STAR METHODS

### Experimental animals

All experiments followed the U.S. National Institutes of Health guidelines for animal research, under an animal use protocol (ARC #2007-035) approved by the Chancellor’s Animal Research Committee and Office for Animal Research Oversight at the University of California, Los Angeles. All mice were housed in a vivarium with a 12/12 h light/dark cycle and experiments were performed during the light cycle The mouse lines used in this study were obtained from Jackson laboratory: *Pv-Cre* mice (JAX 008069), *Ai9*;WT (JAX 007909), *Nkx2.1-Cre;WT* (JAX 008661), *Ai14*;WT (JAX 007914), *Sst-FlpO* (JAX 031629), *Ai65F*;WT (JAX 032864), *Rpl22^HA/HA^* (JAX0110029). Mouse lines were crossed to WT (JAX line 000664) or *Fmr1*^−/−^ female mice (JAX 003025). *Pv-Cre* mice and Ai9 reporter lines (both on C57BL/6J background) were back crossed to FVB WT and *Fmr1*^−/−^ mice for 10-12 generations (Goel et al., 2018). For this study, the following mouse lines were generated: *Nkx2.1-Cre^+/−^*;*Fmr1*^−/−^, Ai14^+/+^;*Fmr1*^−/−^, *Sst-FlpO^+/+^;Fmr1*^−/−^, *Nkx2.1- Cre^+/−^;Sst-FlpO^+/+^*WT, *Nkx2.1-Cre^+/−^;Sst-FlpO^+/+-^*;*Fmr1*^−/−^, *Nkx2.1-Cre^+/−^*;*Rpl22^HA/HA^;WT, Nkx2.1- Cre^+/−^;Rpl22^HA/HA^;Fmr1*^−/−^.

### Viral injections

Depending on the experiment, stereotaxic viral injections were done using glass micropipettes (Sutter Instrument, 1.5 mm outer diameter, 0.86 mm inner diameter) through burr holes (at P0-P1 or P6) or at the time of the cranial window (at P10 or P15; see below). All mice were anesthetized with isoflurane (5% induction, 1-1.5% maintenance via a nose cone, v/v) and placed in a stereotaxic frame with rubber ear bars. Pups were allowed to recover on a warm water circulation blanket before being returned to the dam. For injections at P0-P1, we made a small skin incision on the scalp over the right hemisphere under sterile conditions and drilled a small burr hole over the right S1 cortex using a dental drill, as previously described (C. X. He et al., 2018). A single injection of 250-350 nL of rAAV was done using a Picospritzer (General Valve, 30 pulses of 6 ms, 30 psi). For injections at P10 or P15, we typically pressure-injected 250-350 nL of rAAV at 4-6 different locations over S1 cortex at a depth of 0.2 mm below the dura (30 pulses, 6 ms each, 30 psi). The following viruses were used: AAV1-Syn-GCaMP6s-WPRE-SV40 (Addgene 100843), AAV1-CAG-Flex-GCaMP6s-WPRE.SV40 (Addgene 100842), AAV8-Ef1a-fDIO-GCaMP6s (Addgene 105714), AAV1-hSyn-DIO-hM3D(Gq)-mCherry (Addgene 44361), AAV-hSyn-DIO-mCherry (Addgene 50459-AAV1, 50459-AAV8), AAV8-nEF-Con/Foff-ChRmine-oScarlet, AAV8-EF1a-Con/Foff-2.0-mCherry (Addgene 137133).

### Cranial windows

Pups were anesthetized with isoflurane (5% induction, 1.5–2% maintenance via a nose cone, v/v) and secured in a stereotaxic frame with rubber ear bars. A 3.0–3.5 mm diameter craniotomy was performed over the right S1 cortex, under sterile conditions, by removing the skull without disturbing the pia, as described previously (He et al., 2017; Golshani et al., 2009). The craniotomy was then covered with a 3 mm glass coverslip and secured by cyanoacrylate glue and dental cement. An aluminum head bar was attached to the skull contralaterally to the window with dental cement to secure the animal to the microscope stage. Within 1 h after surgery, the pups appeared fully recovered from the effects of anesthesia and could be returned to their dam. For in vivo imaging at P6 and P10, cranial windows were performed on the day of the recording under 1-1.5% isoflurane anesthesia (5% induction) and we allowed mice to recover for 4-6 h with their dam prior to calcium imaging. For in vivo calcium imaging at P15 or P21, cranial windows were implanted at P10 and P15, respectively, during which we performed GCaMP6s viral injections.

### Intrinsic signal imaging

For animals undergoing calcium imaging at P15 or P21, we previously mapped the location of the barrel field of S1 cortex (S1BF), with intrinsic signal imaging, 1-3 d after cranial window surgery, as described previously (Zeiger et al., 2021). The cortical surface was illuminated by green LEDs (535 nm) to visualize the superficial vasculature. The macroscope was then focused ∼300 μm below the cortical surface and red LEDs (630 nm) were used to record intrinsic signals, with frames collected at 30 Hz 0.9 s before and 1.5 s after stimulation, using a fast CCD camera (Teledyne Dalsa Pantera 1M60), a frame grabber (64 Xcelera-CL PX4, Dalsa), and custom routines written in MATLAB (version 2009a). Thirty trials separated by 20 s were conducted for each imaging session. The whiskers contralateral to the cranial window were bundled together using bone wax and gently attached to a glass microelectrode coupled to a ceramic piezo-actuator (PI127-Physik Instrumente). Each stimulation trial consisted of a 100 Hz sawtooth stimulation lasting 1.5 s. The response signal was divided by the averaged baseline signal, summed for all trials, with a threshold at a fraction (65%) of maximum response to delineate the cortical representation of stimulated whiskers and guide viral injections to S1BF.

### In vivo 2-photon (2P) calcium imaging in head-restrained mice

Calcium imaging was performed on one of two microscopes. For Figs. 1, 5 and Supplementary Fig.9, we used a custom-built 2P microscope and acquired frames (1024×128 down sampled to 256 x 128 pixels, 210×105 µm) at 7.8 fps with a 20x objective (0.95 NA, Olympus) and ScanImage software (Pologruto et al., 2003). For Figs. 2, 4 and Figs. S1, S2, we used a Bergamo II (Thorlabs) at 15.6 fps (512 x 512, or 667 x 667 µm) with the same 20x objective. Both microscopes were coupled to a Chameleon Ultra II Ti:sapphire laser (Coherent) tuned to 930 nm (average power at the sample was <90 mW). We imaged at a depth of 170-190 μm or 220-250 μm below the dura for P6-P10 mice and P15-P21 mice, respectively, corresponding to L2/3. At P15 and P21, mice were lightly sedated with chlorprothixene (2 mg/kg, i.p.) and isoflurane (0.1-0.5%) and kept at 37°C using a temperature control device and heating blanket (Harvard Apparatus). The isoflurane level was manually adjusted to maintain a constant breathing rate (100-150 breaths/min for P15-P16 mice and 140-150 breaths/min for P21 mice). For P6 and P10 experiments mice were unanesthetized and placed in a cotton pad and kept at 37°C using a temperature control device and heating blanket (Harvard Apparatus) (Golshani et al., 2009). Whisker stimulation was achieved by bundling the contralateral whiskers (typically all macrovibrissae of at least 1 cm in length) with soft bone wax and attaching them to a glass capillary coupled to a piezo-actuator.

### In vivo two-photon optogenetic stimulation

Optogenetic photostimulation of ChRmine-expressing neurons (Marshel et al., 2019) was performed by scanning (at 15.1 fps) a separate 1,040 nm femtosecond-pulsed laser (Fidelity, Coherent) using the same 2P microscope (Bergamo II). Laser power was typically 50-60 mW at the objective. Photostimulation consisted of twenty 1 s-long stimulation pulses, with an inter-stimulation interval of 3 s, and was controlled by an optical beam shutter controller (SC10, Thorlabs) triggered via custom-built MATLAB code.

### Nkx2.1-Cre-specific Ribotag RNA extraction and sequencing

Aged matched (P15), *Nkx2.1-Cre^+/−^*;*Rpl22^HA/-^;WT or Nkx2.1-Cre^+/−^;Rpl22^HA/-^;Fmr1*^−/−^, injected at P1 with AAV virus expressing m-Cherry or hM3Dq (see methods and Fig.4) were used for the Ribotag RNA sequencing experiments. For each RNA isolation experiment, coronal sections (1.5 mm thick) containing S1 (A/P -0.5 to +1.82mm, M/L 3.0 to 4.0 mm) were collected and placed in chilled choline-ACSF (132 mM choline, 2.5 mM KCl, 1.25 mM NaH2PO4, 25 mM NaHCO3, 7 mM MgCl2, 0.5 mM CaCl2, and 8 mM D-glucose), from which we dissected S1 bilaterally. Samples were independently homogenized with a 2 mL Dounce homogenizer in 1mL of supplemented homogenization buffer containing: 1% w/v, HB-S; 1 mM DTT, Protease inhibitors (1X), RNAsin (200 units/mL), Cycloheximide (2 mg/l), Heparin (1mg/mL). 100 µL of the homogenate was used for isolating bulk RNA from S1 cortex. To isolate mRNA of HA-tagged Nkx2.1+ IN-specific ribosomes from non-HA tagged ribosomes, samples were incubated for 4 h with an anti-HA antibody (1:180, 5 µL in 900 µL of homogenate; 901513 Covance Anti-HA antibody), followed by overnight incubation with magnetic beads (Pierce Protein A/G Magnetic Beads 88802, 200 µL) at 4°C at 15 rpm centrifugation. The supernatant was washed thrice with 800 μL of high salt buffer (Tris 50mM, KCl 300mM, MgCl_2_ 12mM, 10% NP-40 (1%), DTT 1mM, Cycloheximide 100μg/ml, dH_2_O), centrifuged at 15 rpm for 10 min at 4°C. For RNA extraction, we used Zymo Research Direct-zol RNA Miniprep kit (R2051) for the homogenate (input), and the Zymo Research Direct-zol RNA Microprep kit (R2061) for the bead-antibody-protein sample (IP); RNA was eluted with DNAase/RNAse-free water. RNA Integrity Number (RIN) scores were used to evaluate the integrity of RNA through the ratio of 28S:18S ribosomal RNA (Agilent TapeStation 4200) (Schroeder et al., 2006). We set a minimum of RIN score 7 for our samples (Gandal et al., 2018). RNA sequencing libraries were prepared using Ovation® RNA-Seq System V2. Libraries were multiplexed for paired-end 50 bp sequencing on NovaSeq S2, with read depth of 50 million reads on average. Demultiplex reads were aligned using STAR to the mouse genome (reference genome ID: mm10) and fragment counts were derived using HTS-seq (Anders et al., 2015). Outliers were calculated by measuring connectivity between samples, using the WCGNA package (Langfelder and Horvath, 2008) and removing samples >2 SD from the mean for Bulk and IP samples. All analysis was performed using R 4.0.3. Differential expression analysis was performed using the Bioconductor DESEQ2 package (Love et al., 2014) with FDR of 0.1. Go enrichment analysis was performed using Bioconductor clusterProfiler 4.0 (Wu et al., 2021).

### Data analysis for calcium imaging

Calcium imaging data were analyzed using custom-written MATLAB routines (MATLAB version 2020a). *X*–*Y* drift in the movies was first corrected using either a frame-by-frame, hidden Markov model-based registration routine (Dombeck et al., 2007) or a cross-correlation-based, nonrigid alignment algorithm (Mineault et al., 2016). The choice of registration algorithm did not affect the data analysis since the fluorescence data for each neuron was always normalized to its own baseline. For ROI segmentation, we used either EZcalcium (Cantu et al., 2020) (for Figs. 1, 5D-G and Supplementary data Fig.9) or Suite2P (Pachitariu et al., 2016) (for Figs. 2, 4I-M, and Supplementary Fig.2). To quantify activity levels, a “modified *Z* score” *Z*_*F* vector (an epoch was 8 consecutive frames wherein *Z*_*F>* 3) for each neuron was calculated as *Z*_*F* [*F*(*t*) mean(quietest period)]/SD(quietest period), where the quietest period is the 10 s period with the lowest variation (standard deviation) in d*F*/*F*, as described previously (C. X. He et al., 2017). All subsequent analyses were performed using the *Z*_*F* vectors. To define whether an individual cell showed responses that coincided with epochs of whisker stimulation (“stimulus-locked”), a probabilistic bootstrapping (10,000 scrambles) method was implemented as previously described (C. X. He et al., 2017). To assess adaptation of neuronal activity to repetitive whisker stimulation we calculated an adaptation index: [(*Z* score during first five stimulations) > (*Z* score during last five stimulations)]/[(*Z* score during first five stimulations) > (*Z* score during last five stimulations)]. The percentage of active cells was based on the Z_F vector, calculated as of proportion of cells reaching the criterion of having at least one activity epoch with Z_F*>* 3 for 8 consecutive frames, amongst the total number of segmented ROIs. A Pearson’s correlation coefficient was calculated for all neuron pairs in each calcium imaging movie using the Z_F vector.

### Brain slice electrophysiology

*Pv*-Cre^+^;tdTom^+^;*Fmr1^−/−^*mice at P15 were deeply anesthetized using 5% isoflurane prior to decapitation and 400 µm-thick coronal slices were prepared with a vibratome (Leica VT1200S, Germany). Slices were prepared in ice-cold oxygenated (95% O_2_ and 5% CO_2_) modified artificial cerebrospinal fluid (ACSF) containing (in mM): 92 NMDG, 5 Na^+^-ascorbate, 3 Na^+^-Pyruvate 2.5 KCl, 10 MgSO_4_, 2 CaCl_2_, 1.2 Na2HPO_4_, 24 NaHCO_3_, 5 HEPES, 25 Glucose. Slices were then left to recover in the modified ACSF at ∼33°C for 30 min before being transferred to oxygenated regular ACSF at room temperature (RT) for storage for at least 1 h prior to recording. Recordings were made at 32-33°C in oxygenated ACSF containing (in mM) 124 NaCl, 2.5 KCl, 2 MgSO_4_, 2 CaCl_2_, 1.2 Na2HPO_4_, 24 NaHCO_3_, 5 HEPES, 13 Glucose, and the perfusion rate was set to 5 mL/min. Whole-cell patch-clamp recordings were performed in cortical L2/3 *Pv*-Cre^+/−^-tdTom^+^ cells. Recordings relied on fluorescence visualization using a Zeiss AxioSkop FS+ microscope and additional verification by their intrinsic electrophysiological properties. For current-clamp recordings, the intracellular solution consisted of (in mM) 100 K-gluconate, 20 KCl, 4 ATP-Mg, 10 phosphocreatine, 0.3 GTP-Na, 10 HEPES (adjusted to pH 7.3 and 300 mOsm). Intrinsic excitability was measured as the number of action potentials evoked during a 250 ms current step at intensities of 0.05, 0.1, 0.15, 0.2, 0.25, 0.3, 0.35, 0.4, 0.45, 0.5 nA. Wash-on of AG00563 (10 µM) was performed at 32-33°C for a minimum of 5 min before assessing PV-IN intrinsic excitability. Series resistance was monitored, and recordings were discarded if series resistance changed by more than 15%, or if apparent loss of current clamp control occurred as reflected by a sudden change in the recording stability.

### Immunohistochemistry and IN quantification in mouse tissue

Mice were anesthetized with 5% isoflurane and transcardially perfused with ice cold phosphate buffer saline PBS (0.1 M) followed by ice cold 4% paraformaldehyde in PBS and post-fixed overnight at 4°C. Coronal sections (60 μm) were cut on a vibratome (Leica VT1200S). Sections were permeabilized with 0.3% Triton X-100 and blocked with 5% normal goat serum (NGS) for 1 h at RT. Sections were then incubated overnight at 4°C with the primary antibody diluted in PBS with 5% NGS and 0.1% Triton X-100. After rinsing in PBS for 5-10 min 3 times, sections were incubated for 1 h at RT with the corresponding Alexa Fluor-conjugated secondary antibody diluted in PBS (1:1000, Invitrogen). After rinsing in PBS for 5-10 min 3 times, sections were mounted onto Superfrost Plus glass slides (Vectashield Vibrance) with DAPI mounting medium and stored in the dark at RT. The following primary antibodies were used: mouse anti-parvalbumin (1:1,000, Sigma SAB4200545), rat anti-somatostatin (1:300, ab108456, Abcam), goat anti-mCherry (1:1000, M11240, Millipore), rabbit anti cleaved-Caspase-3 (1:400, D175-5A1E, Cell Signalling).

To quantify IN density, sections were imaged on an ApoTome2 microscope (Zen2 software, Zeiss; 10X objective, 0.3 NA), and analyzed with ImageJ (https://imagej.nih.gov/ij/). The section sampling fraction (ssf=1/6) for S1BF was determined within [A/P: bregma -0.94mm to -1.94mm]. Composite images spanning the S1BF were acquired by stitching a grid of 4-6 confocal images (2754×2061 pixels). For Nkx2.1Cre^+^;tdTom^+^ and Pv-Cre^+^;tdTom^+^ cell count was performed by automated detection using ImageJ plug-ins (Filter minima=1.5 pixel, Process maxima prominence>200pixels). PVALB^+^ and Cleaved-Caspase3^+^ cell count was performed using Cell counter plug-in. Cortical layers were determined using DAPI counterstaining and cell counts were subdivided into layers 1-3, 4 and 5-6. The size of the total cortical area and layer-specific ROIs were measured using ImageJ. Cell counts were divided by ROI area and reported as cells/mm^2^.

### Histology and IN quantification in human tissue

We collected post-mortem cortical samples of Brodmann Area 3 from 8 FXS cases and 8 neurotypical, sex and age-matched cases (**Table 1**). The formalin-fixed tissue was obtained from CENE (Hispano-American brain bank of Neurodevelopmental Conditions, https://health.ucdavis.edu/mindinstitute/research/cene-brain-bank/cene-about.html). The neurotypical cases were determined to be free of neurological disorders, including autism or intellectual disability, based on medical records and information gathered at the time of death from next of kin. All cases were males, except for one control case and one FXS case. Tissue blocks were fixed in 10% buffered formalin, cryoprotected in a 30% sucrose solution in 0.1 M phosphate-buffered saline with 0.1% sodium azide. Tissue was embedded in optimum cutting temperature compound, and frozen at −80 °C. A cryostat was used to cut 14 μm-thick slide-mounted sections, stored at −80 °C until use. Based on Brodmann cortical neuroanatomy, blocks containing area BA3 (primary somatosensory cortex) were isolated from each case. We cut 14 µm sections on a cryostat, stained 1 section with Nissl, and chose sections that exactly matched the von Economo descriptions for BA3. On adjacent sections we performed triple immunostaining for PV, Calbindin (CB), and Calretinin (CR). We used the following primary antibodies for our quantification experiments: monoclonal mouse anti-CB-D28k (1:500, Swant 300, Switzerland), polyclonal rabbit anti-CR (1:500, Swant 7697), and monoclonal mouse anti-PV (1:500, Swant 235). The following primary antibodies were used for our validation study: polyclonal rabbit anti-CB (1:500, Abcam ab11426), monoclonal mouse anti-CR (1:500, Millipore mab1568), and polyclonal rabbit anti-PV (1:500, Abcam ab11427). Secondary antibodies were donkey anti-mouse conjugated with biotin, amplified with avidin-biotin complex (ABC) and developed with diaminobenzidine (DAB) or Vector NovaRED substrates (all from Vector, USA) for CB and PV detection, respectively. Donkey anti-rabbit antibody conjugated with alkaline phosphatase and Vector Blue substrate (Vector) was used for CR detection. Tissue was pretreated in Diva decloaker (DV2004 LX, MX, Biocare medical, USA) in a decloaking chamber (Biocare medical, USA) at 110°C for 6 min. Subsequently, immunostained tissues were immersed in 0.1% Nissl for 1 min and then dehydrated in successive baths of 50% ethanol (30 s), 70% ethanol (30 s), 95% ethanol (10 min), and 100% ethanol (5 min), isopropanol (5 min), followed by xylene (15 min). Sections were then mounted and coverslipped with Permount (Fisher). The PV-red staining was visualized as pink following Nissl staining. We quantified the number of immunopositive cells for each interneuron subtype (CB+, CR+, or PV+ cells). We processed 1 section of each case and chose a 3 mm wide bin parallel to the pial surface that extended through the cortical gray matter to include all cortical layers. We imaged the tissue a 100X oil objective (Olympus BX61 microscope Hamamatsu Camera) and quantified IN density using MBF Bioscience StereoInvestigator V.9 Software (MicroBrightField, Williston, VT), on a Precision PWS 690, Intel Xeon CPU computer (Dell) with Windows XP Professional V.2002 system (Microsoft).

### DREADD agonist (C21) administration in vivo

For DREADD experiments, the agonist C21 (HelloBio) was dissolved in 0.9% saline to 2 mg/mL. For acute DREADD experiments at P15 in *Pv-Cre;Fmr1^−/−^* mice, we injected C21 at 1 mg/kg, s.c. (Thompson et al., 2018). For the chronic DREADDs experiments in Nkx2.1 pups we injected C21 twice daily from P5 to P9 at a dose of 1 mg/kg, s.c.

### AG00563 treatment in vivo

AG00563 (1-(4-methylbenzene-1-sulfonyl)-N-[(1,3-oxazol-2-yl)methyl]-1H-pyrrole-3-carboxamide) was synthesized by Lundbeck. Purified AG00563 was shipped to UCLA for reconstitution and experiments. AG00563 was dissolved in a solution of 10% of hydroxypropyl-b-cyclodextrine (Sigma) in 0.9% saline. For the Kv3.1 pharmacological manipulation at P21, mice were injected with AG000563 (3 mg/kg, s.c.) either once acutely at P15, or chronically (twice daily) from P15 to P20. Control animals were injected with a vehicle solution containing 10% of hydroxypropyl-beta cyclodextrine in 0.9% saline.

### Tactile defensiveness assay in head-restrained mice

We used a paradigm we previously established to assess tactile defensiveness in juvenile and adult mice (C. X. He et al., 2017). Briefly, titanium head bars were implanted at P15 and then mice were habituated to head restraint and to running on an air-suspended 200 mm polystyrene ball. For habituation, mice were placed on the ball for 20 min/d for 5 consecutive days before testing at P21. On the test day, each animal was first placed on the ball for a 3 min baseline period. Next, we performed a sham stimulation trial in which the whisker stimulator was visibly moving, but just out of the reach of the whiskers on the animal’s left side. The stimulator consisted of a long, narrow comb of five slightly flexible von Frey Nylon filaments that were attached to a piezoelectric actuator (Physyk Instrumente). During the stimulation trial, the filaments were intercalated between the whiskers. Bundling of whiskers onto a glass capillary (as was done for calcium imaging) was not feasible for these awake experiments because the mouse could have damaged the capillary or unbundled some of its whiskers with its forepaw. The stimulation protocol consisted of a 10 s baseline followed by 20 sequential bouts of whisker deflections along the anterior–posterior direction (1 s long at 10 Hz), with a 3 s interstimulus interval, and ending with another 10 s post-stimulation baseline. A custom-written semiautomated video analysis was implemented in MATLAB to score defensive behaviors (grabbing the stimulator) and adaptive healthy behaviors (grooming), in each 1s increment of the videos during the 20 stimulations (C. X. He et al., 2017).

### Statistical analyses

Unless otherwise specified, results were plotted and tested for statistical significance using Prism 9. Statistical analyses of normality (Lilliefors and Shapiro Wilk tests) were performed on each data set; depending on whether the data significantly deviated from normality (p<0.05) or not (p>0.05), appropriate non-parametric or parametric tests were performed. The statistical tests performed are mentioned in the text and the legends. For parametric two-group analyses, a Student’s *t*-test (paired or unpaired) was used. For non-parametric tests, we used Mann-Whitney test (two groups) and the Friedman test (repeated measures). Multiple comparisons with single variables were analyzed using one-way ANOVA with post-hoc Bonferroni’s test (comparing the mean of each column with the mean of every other column) or Dunnett’s test (comparing the mean of each column with the mean of a control column) for normally distributed samples. For multiple comparisons with more than one variable, a two-way ANOVA with post hoc Sidak’s test was used. No statistical methods were used to predetermine sample size. Sample sizes were calculated based on similar published studies. In the figures, significance levels are represented with the following convention: * for p<0.05; ** for p<0.01, *** for p<0.001. All experiments were replicated at least in 3 different litters. In all the figures, we plot the standard error of the mean (s.e.m.). Graphs either show individual data points from each animal or group means (averaged over different mice) superimposed on individual data points.

---

**Supplementary Fig.1:**
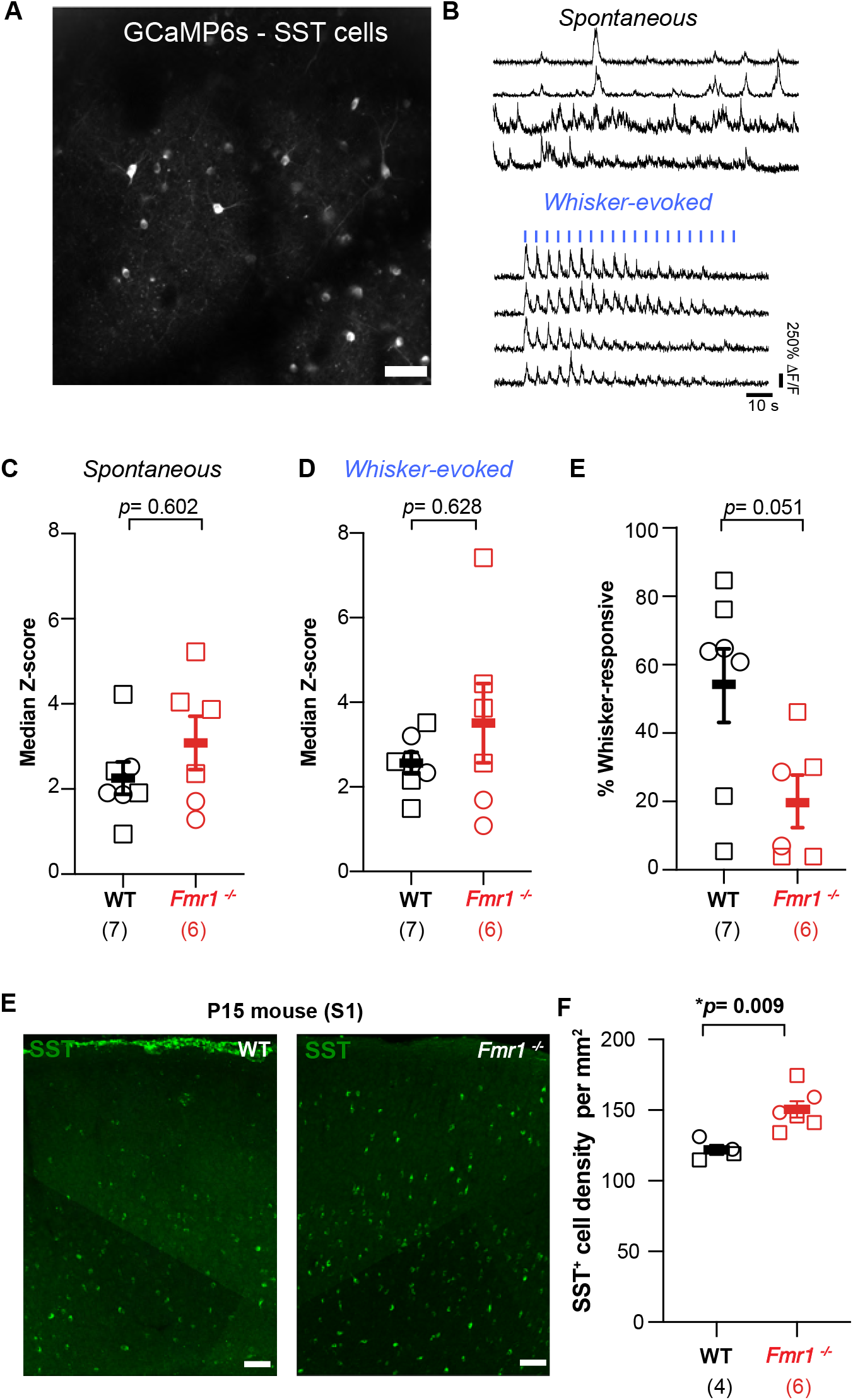
SST activity is unremarkable in S1 of P15 *Fmr1*^−/−^ mice. a. Example field of view of SST-INs expressing GCaMP6s in SST-Flp mice at P15. b. Example traces of calcium transients during spontaneous and whisker-evoked recordings. in both WT and *Fmr1*^−/−^ mice (traces are from 2 different SST-INs in two different animals). The vertical blue bars represent the 20 whisker stimulations. c. Mean Z-scores for spontaneous activity of SST-INs in *Fmr1*^−/−^ and WT mice at P15 (2.56 ± 0.38 for WT vs. 3.08 ± 0.62 for *Fmr1*^−/−^, n=6 and 7 mice, respectively; p=0.602, MW t-test). d. Mean Z-scores for whisker-evoked activity of SST-INs in *Fmr1*^−/−^ and WT mice at P15 (2.56 ± 0.25 for WT vs. 3.51 ± 0.94 for *Fmr1*^−/−^, p=0.628, MW t-test). e. Percentage of whisker-responsive SST-INs in *Fmr1*^−/−^ and WT mice at P15 (53.8 ± 11.0% for WT vs. 19.6 ± 7.6% for *Fmr1*^−/−^, p= 0.051, M-W t-test). f. Coronal section through the barrel field of S1 in WT and *Fmr1*^−/−^ mice immunostained for SST (green) across the entire cortical thickness. Scale bar= 50µm g. Quantification of total SST^+^ cell density in the barrel field of S1 in WT and *Fmr1*^−/−^ mice at P15 (WT: 121.9 ± 3.7 cells/mm^2^, *Fmr1*^−/−^: 150.5 ± 5.9, n= 4 and 6, respectively; p=0.009, MW test).

**Supplementary Fig.2:**
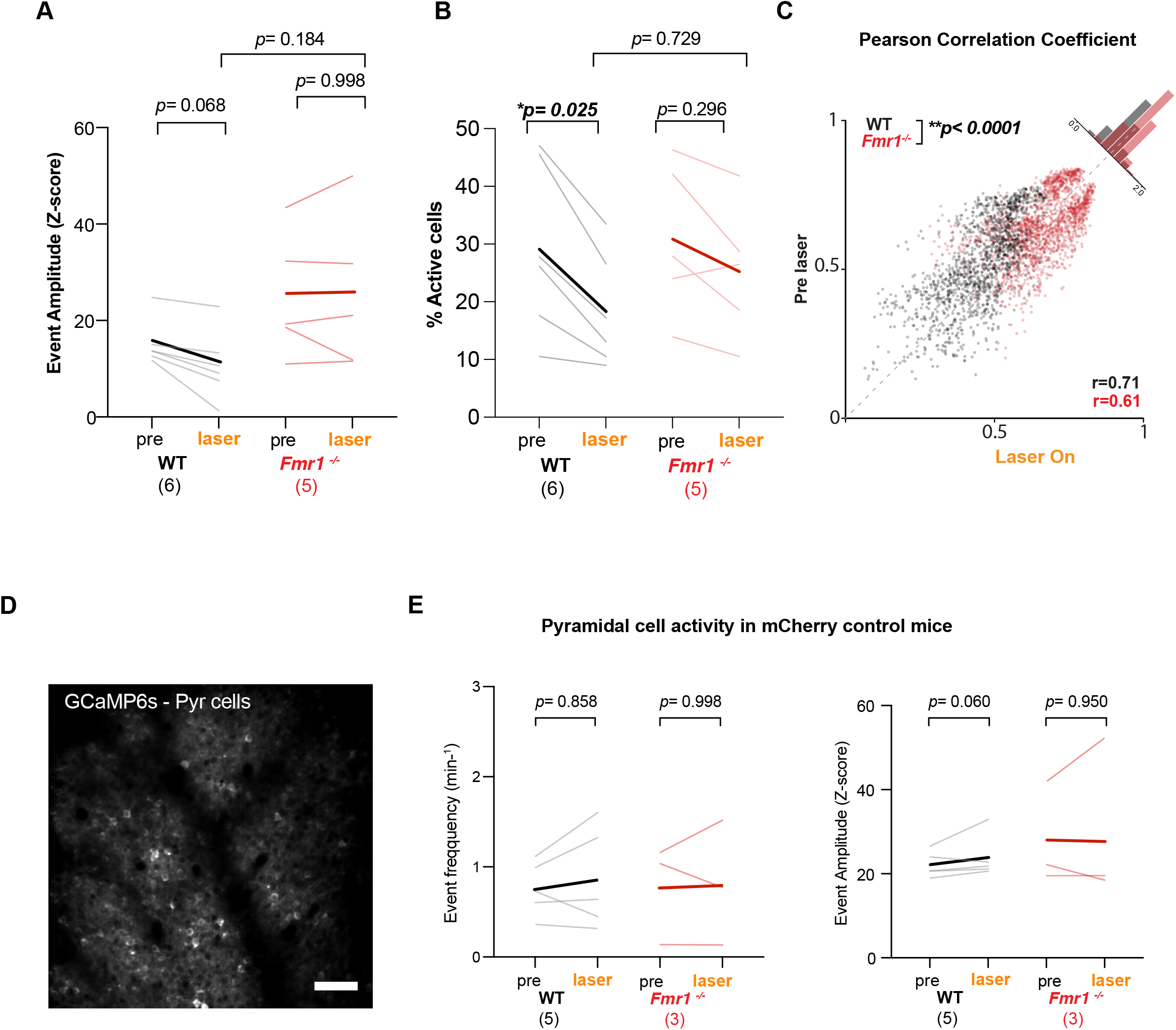
Optogenetic stimulation of Nkx2.1-INs fails to modulate Pyr neurons in *Fmr1^−/−^* mice and has no effect in mCherry controls (no opsin). a. Amplitude of synchronous calcium events in *Nkx2.1-Cre^+/−^;Sst-FlpO^+/−^* in WT and *Fmr1*^−/−^ mice (n=5 and 6, respectively) after optogenetic activation of MGE-INs (15.3 ± 2.0 pre laser vs. 10. 8 ± 2.9 laser On, p=0.068 in WT mice; 24.9 ± 5.8 pre laser vs. 25.3 ± 7.2 laser On, p=0.998, two-way ANOVA post-hoc Tukey). b. The percentage of active Pyr cells is significantly reduced in WT upon laser stimulation but is unchanged in *Fmr1^−/−^* mice (29.1 ± 6.0 pre laser vs. 18.3 ± 4.0 laser On, p=0.025 in WT mice and 30.9 ± 5.9 pre laser vs. 25.2 ± 5.2 laser On, p=0.296, two-way ANOVA post-hoc Tukey). c. Change in pairwise correlation coefficients after optogenetic MGE-IN activation in *Nkx2.1-Cre^+/−^;Sst-FlpO^+/−^* mice for individual pairs of neurons (Spearman r=0.71, n=1,449 neuron pairs from n=6 WT mice; Spearman r=.0.61, n=1,842 neuron pairs from n=5 *Fmr1*^−/−.^mice). Note that correlation coefficients are significantly reduced by laser stimulation in WT mice, but not in *Fmr1*^−/−^ mice, as shown by the frequency distribution (p<0.001, Kolmogorov-Smirnov test). d. Example field of view of Pyr neurons expressing GCaMP6s in S1 at P10 in a control *Nkx2.1-Cre^+/−^;Sst-FlpO^+/−^* WT mouse expressing mCherry alone (no ChRmine). e. Mean frequency of Pyr cell calcium transients remained unchanged upon optogenetic stimulation in *Nkx2.1-Cre^+/−^;Sst-FlpO^+/−^* WT and *Fmr1*^−/−^ mice that do not express the opsin ChRmine (0.67 ± 0.13 pre laser vs. 0.86 ± 0.25 laser On, p=0.312 in WT mice; 0.78 ± 0.32 pre laser vs. 0.81 ± 0.4 laser On, p=0.500, n=5 WT and n=3 *Fmr1*^−/−^, two-way ANOVA, post-hoc Tukey). f. Mean amplitude of Pyr cell calcium transients remained unchanged upon optogenetic stimulation in *Nkx2.1-Cre^+/−^;Sst-FlpO^+/−^* WT and *Fmr1*^−/−^ mice that do not express the opsin ChRmine (22.18 ± 1.37 pre laser vs. 23.86 ± 2.29 laser On, p=0.156 in WT mice; 0.78 ± 0.32 pre laser vs. 0.81 ± 0.4 laser On, p=0.375, n=5 WT and n=3 *Fmr1*^−/−^ Wilcoxon matched-paired signed rank test).

**Supplementary Fig.3:**
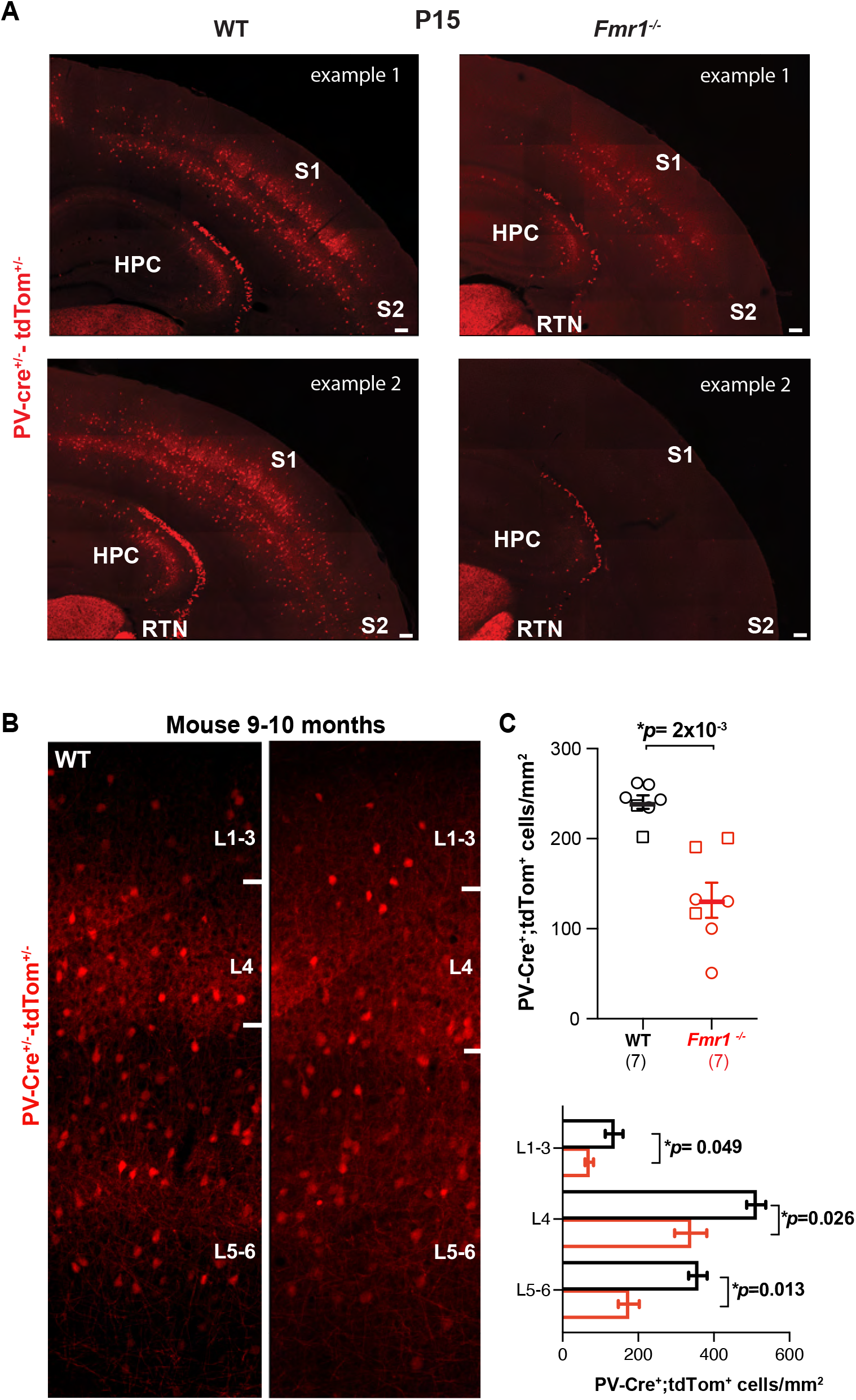
Reduced PV-IN density in *Fmr1*^−/−^ mice at P15 and 9-10 months. a. Example coronal sections through S1 from *PV*-Cre;tdTom mice (WT and *Fmr1^−/−^*) at P15 showing the range of PV-IN density in *Fmr1^−/−^* mice across the dorsal brain. Notably some *Fmr1^−/−^* mice (example 2) exhibit a dramatic loss of PV-INs in neocortex and hippocampus (HPC), while other bran regions, such as the reticular thalamic nucleus (RTN) are much less affected. S2: secondary somatosensory cortex. Scale bars: 100 μm. b. Coronal sections through the barrel field of S1 from *PV*-Cre;tdTom mice (WT and *Fmr1^−/−^*) at 9-10 months (corresponding approximately to age of human tissue in Fig. 3E-F). Scale bar: 50 μm. c. Mean density of PV-tdTom*+* INs in S1 is significantly lower in adult *Fmr1^−/−^* mice (top), even across individual cortical layers (bottom). (all layers: 241 ± 8 cells/mm^2^ for WT vs. 132 ± 2 for *Fmr1*^−/−^; p=0.002, MW *t*-test; L2/3: 136 ± 23 vs. 69 ± 10, p=0.049, L4: 512 ± 26 vs. 339 ± 43, p=0.026; L5/6: 358 ± 24, two-way ANOVA, post-hoc Holm-Sidak test, p=0.013, n=7 per genotype).

**Supplementary Fig.4:**
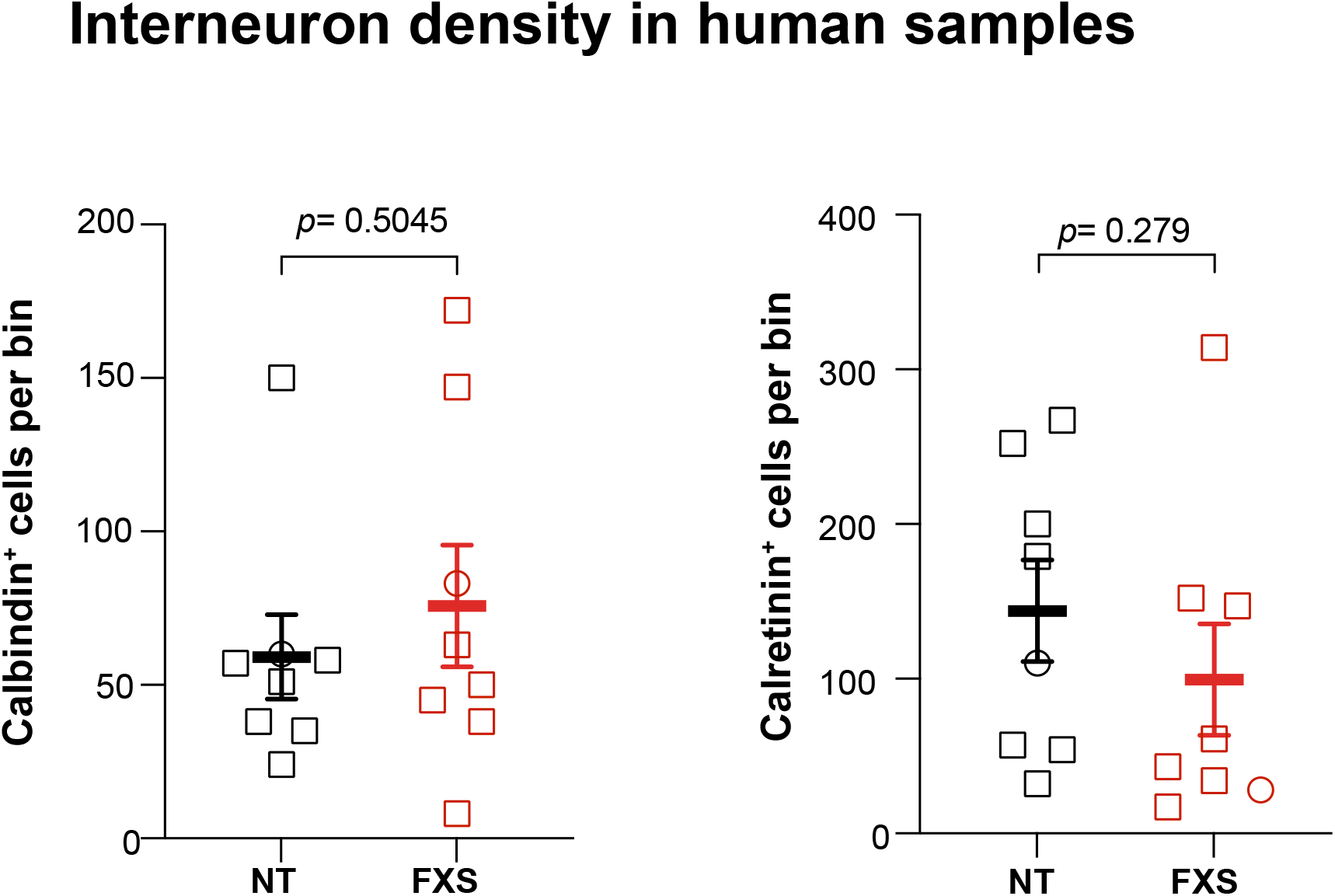
Density of Calbindin- and Calretinin-immunoreactive INs is unchanged in BA3 of FXS human cases. a. Quantification of Calretinin-immunoreactive INs in FXS cases and neurotypical (NT) controls (n=8 per group, 144 ± 33 for NT vs. 100 ± 36 for FXS, p=0.279 MW *t* test). b. Quantification of Calbindin-immunoreactive INs in FXS and NT cases (n=8 per group, 59 ± 14 for NT vs. 76 ± 20 for FXS, p=0.279 MW *t* test).

**Supplementary Fig.5:**
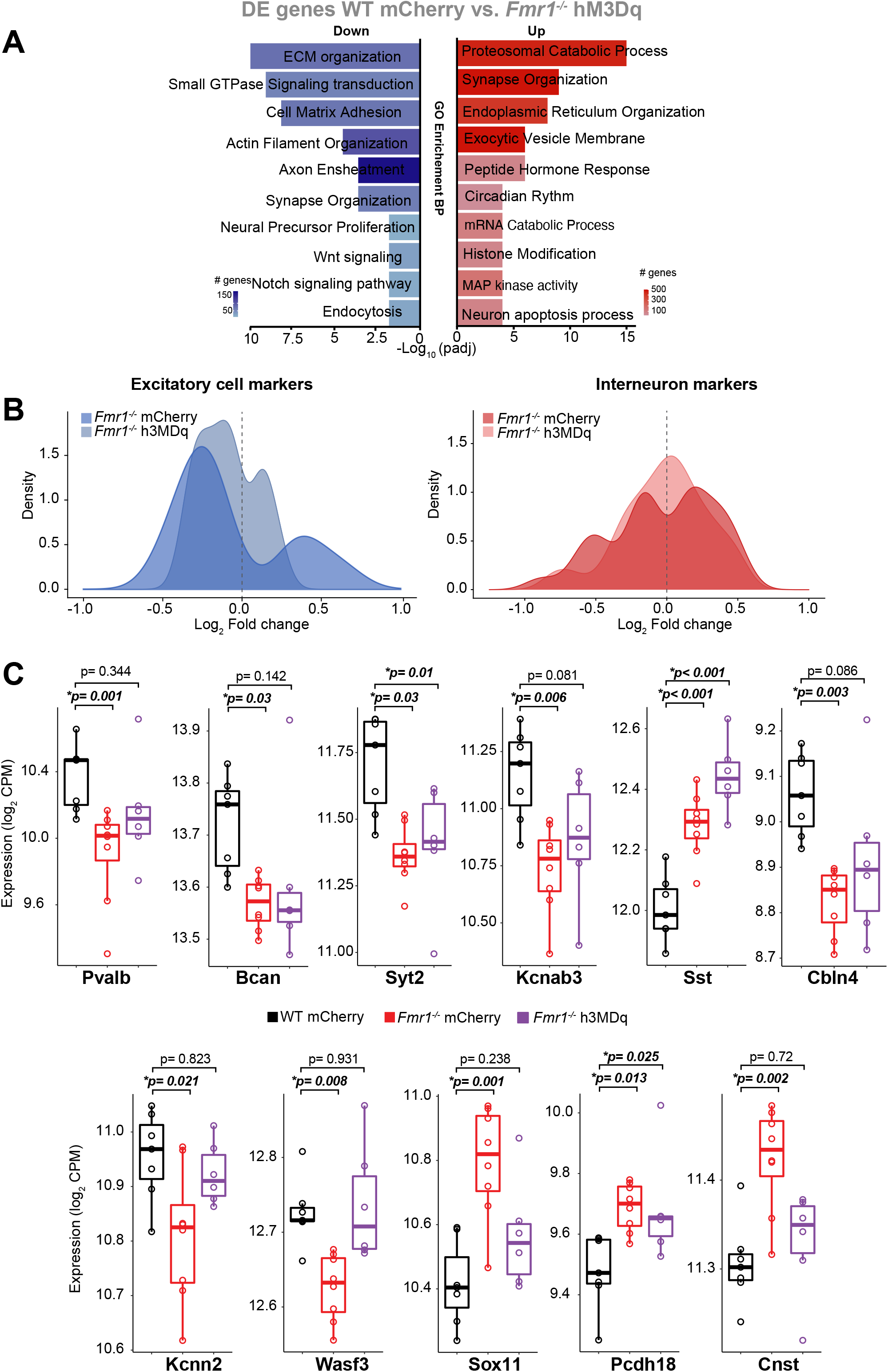
Gene ontology (GO) analysis for the DE genes in *Fmr1*^−/−^-hM3Dq mice compared to WT-mCherry mice. a. GO terms enriched among downregulated (blue) and upregulated (red) genes using the biological process package (BP) (right panel). Scale bars represent the number of genes in each category. Note that “*Synapse organization*” and “*Neuron* a*poptotic process*” categories are less different than in the comparison shown in Fig. 4C between *Fmr1^−/−^* (mCherry) and WT (mCherry) mice, suggesting they were ‘improved’ by the chemogenetic activation of MGE-INs. b. Density plot showing how the log_2_ fold change was affected by the DREADD manipulation in excitatory cells (Tasic et al., 2018) and the interneuron population (Favuzzi et al., 2019; Mahadevan et al., 2021; Paul et al., 2017). Note that fewer genes are different from WT mice in the *Fmr1*^−/−^-hM3Dq group. c. Whisker plots comparing the expression of several MGE-derived markers (PV-INs, including fast-spiking basket cells and chandelier cells, and SST-INs markers; Wilcoxon signed-rank test)

**Supplementary Fig.6:**
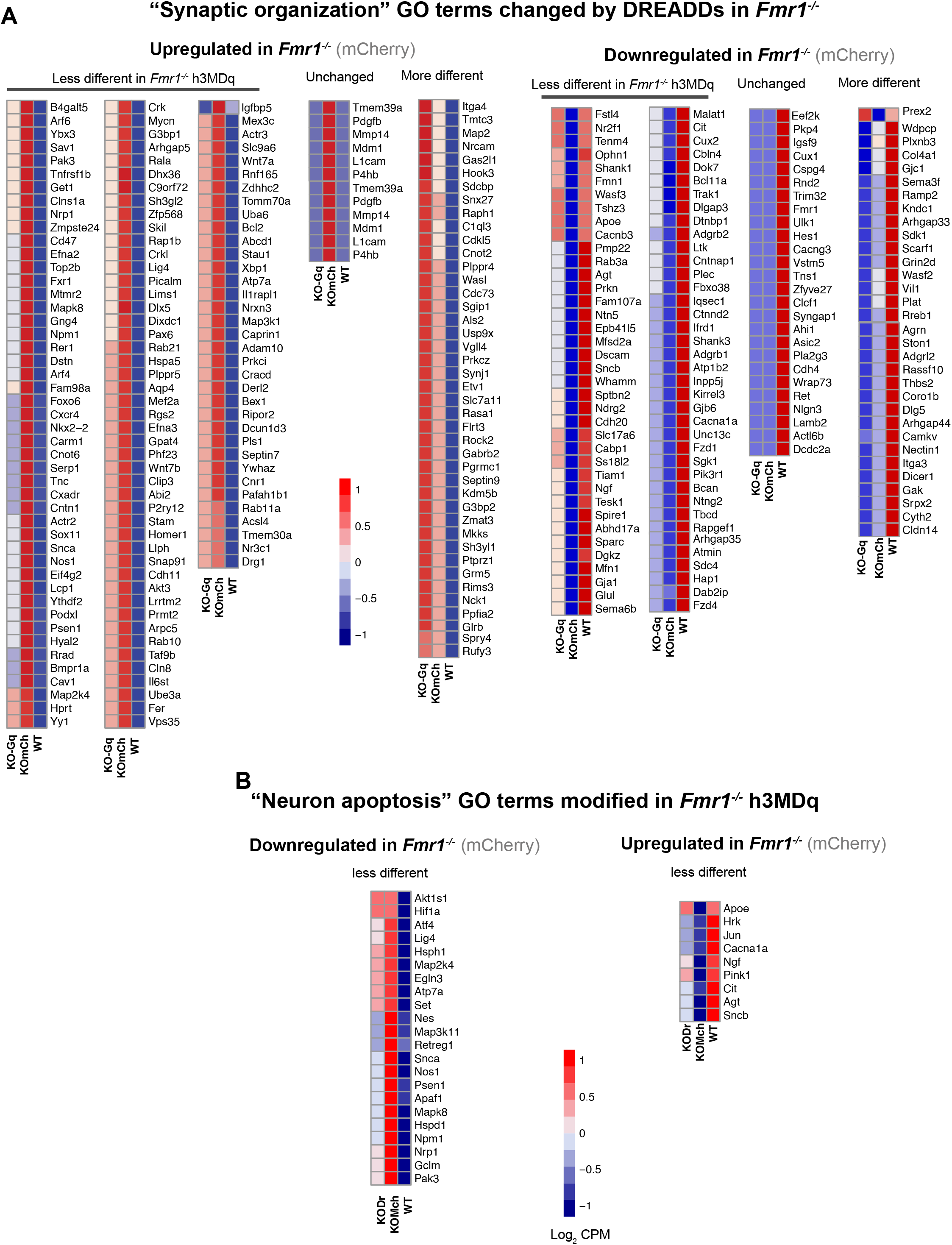
Gene expression levels changed by C21 treatment in *Fmr1*^−/−^-hM3Dq mice. a. List of upregulated or downregulated genes among the GO term “*Synapse organization*” in *Fmr1^−/−^*-mCherry group (as compared to WT-mCherry) that were changed by chronic C21 injections in *Fmr1^−/−^*-hm3Dq mice. Heatmaps represent the average for each treatment/genotype group in (log_2_ CPM). Note that, while DREADD treatment reduced differences for the vast majority of genes. The expression of a few other genes was either unhanged or even more different after C21 treatment. b. Same as in panel A but for Go term “*Neuron* a*poptotic process*.”

**Supplementary Fig.7:**
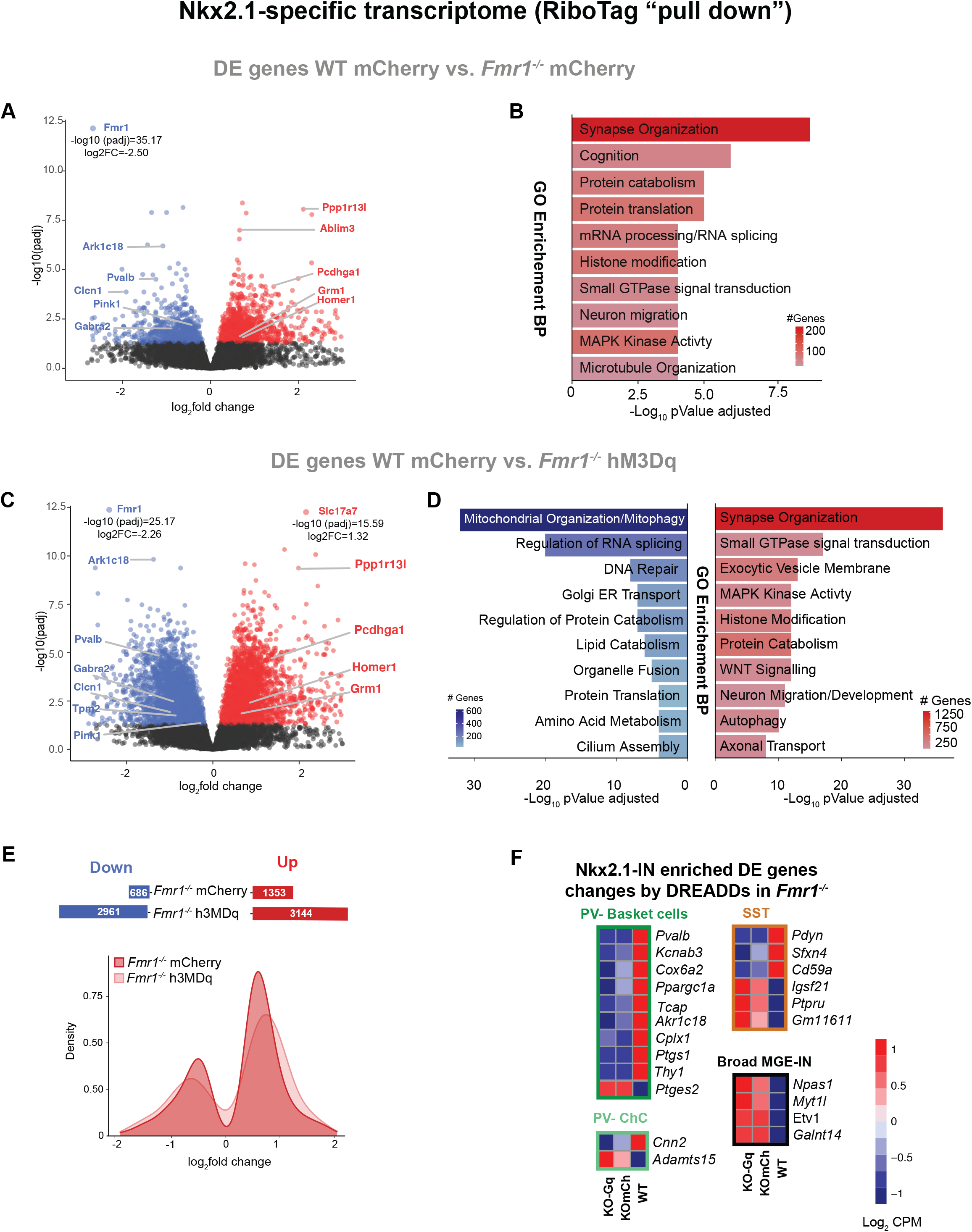
Nkx2.1Cre^+^-IN specific transcriptome. a. Volcano plot of differentially expressed (DE) genes in the pulldown RNA of *Fmr1*^−/−^-mCherry mice (n=8) compared to WT-mCherry control group (n=3). A few genes are annotated (because of their role in IN and synapse function) amongst significantly downregulated (blue) or upregulated genes (red). b. Top 10 ‘GO terms’ using the biological process package for DE genes in *Fmr1*^−/−^-mCherry mice compared to WT-mCherry control group (see Methods). c. Volcano plot of DE genes in the pulldown RNA of *Fmr1*^−/−^-hM3Dq mice (n=6) compared to WT-mCherry controls. d. Top 10 ‘GO terms’ using the biological process package for differentially expressed genes in *Fmr1*^−/−^-hM3Dq mice compared to WT-mCherry control group (see Methods). e. Top: The total number of down- and up-regulated genes (compared to WT-mCherry controls) was higher in *Fmr1*^−/−^-hM3Dq mice than in *Fmr1*^−/−^-mCherry mice. Bottom: Density plot showing how the log_2_ fold change was affected by the DREADD manipulation (more genes are different from WT mice in the *Fmr1*^−/−^-hM3Dq group). f. List of genes associated with different subclasses of MGE-INs for which expression was changed by DREADDs (dark green: basket cells; light green: Chandelier cells; brown: SST cells; black: global MGE-IN markers. Scale bar represents the log_2_ CPM (count per millions).

**Supplementary Fig.8:**
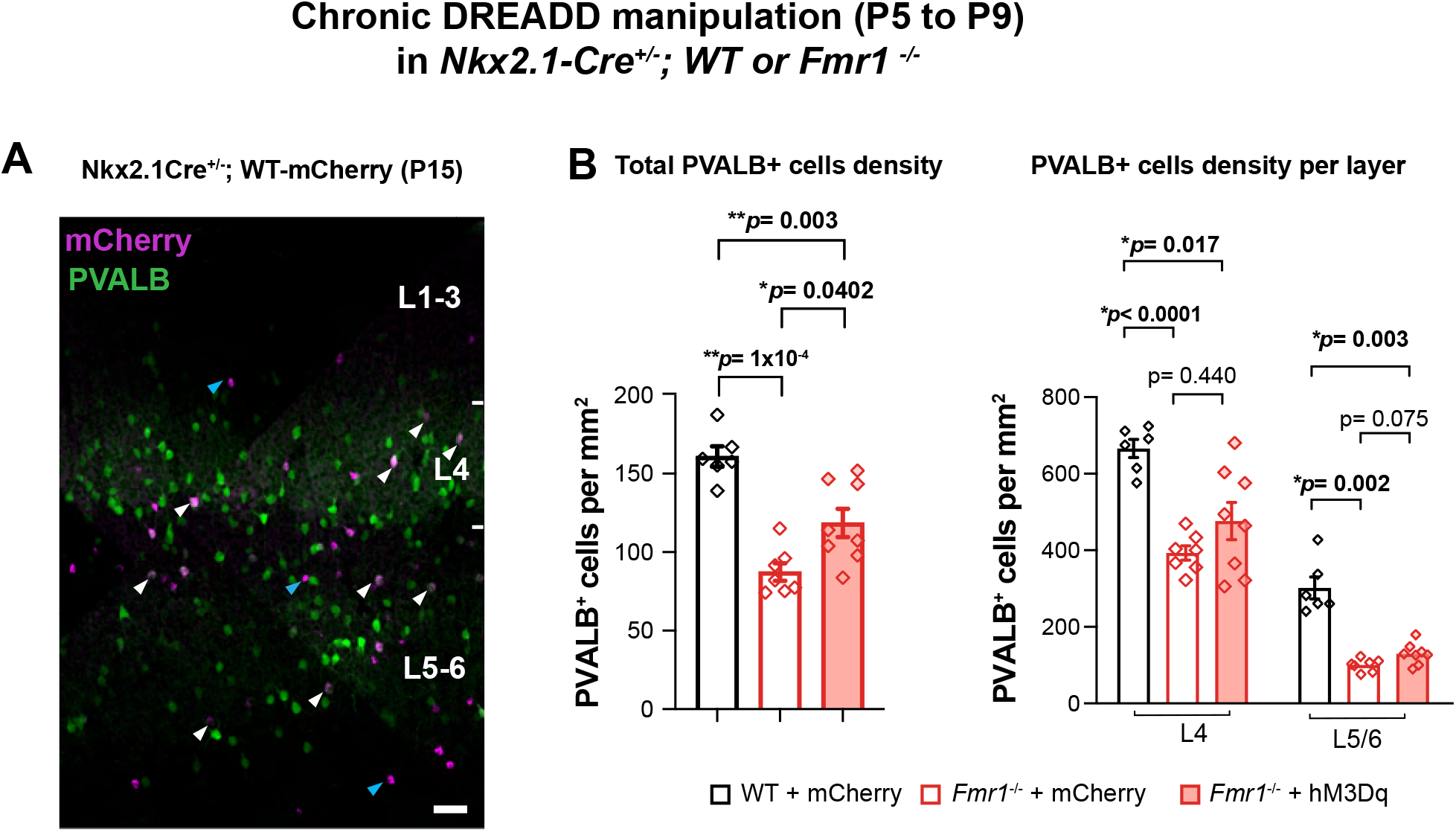
Chronic chemogenetic activation of Nkx2.1-INs (P5-P9) increases the density of PVALB+ cells at P21. a. Coronal section through the barrel field of S1 in a P15 *Nkx2.1-Cre*;mCherry WT mouse. Note the overlap in expression of mCherry (purple) in Nkx2.1+ cells and PVALB immunoreactivity (green) across cortical layers (white arrowheads). As expected, some of the mCherry+ INs do not express PVALB (blue arrowheads); this population includes SST-INs. Scale bar= 50µm. b. Left, Quantification of PVALB+-INs density across cortical layers at P15. (161 ± 6 cells/mm^2^ for WT-mCherry vs. 87 ± 6 for *Fmr1*^−/−^-mCherry, p=0.003; and 119 ± 9 for *Fmr1*^−/−^-h3MDq, p= 0.040, n=6, 7 and 8, respectively). Right, The chronic DREADD manipulation did not significantly change the density of PV-INs in L4 or L5/6.

**Supplementary Fig.9:**
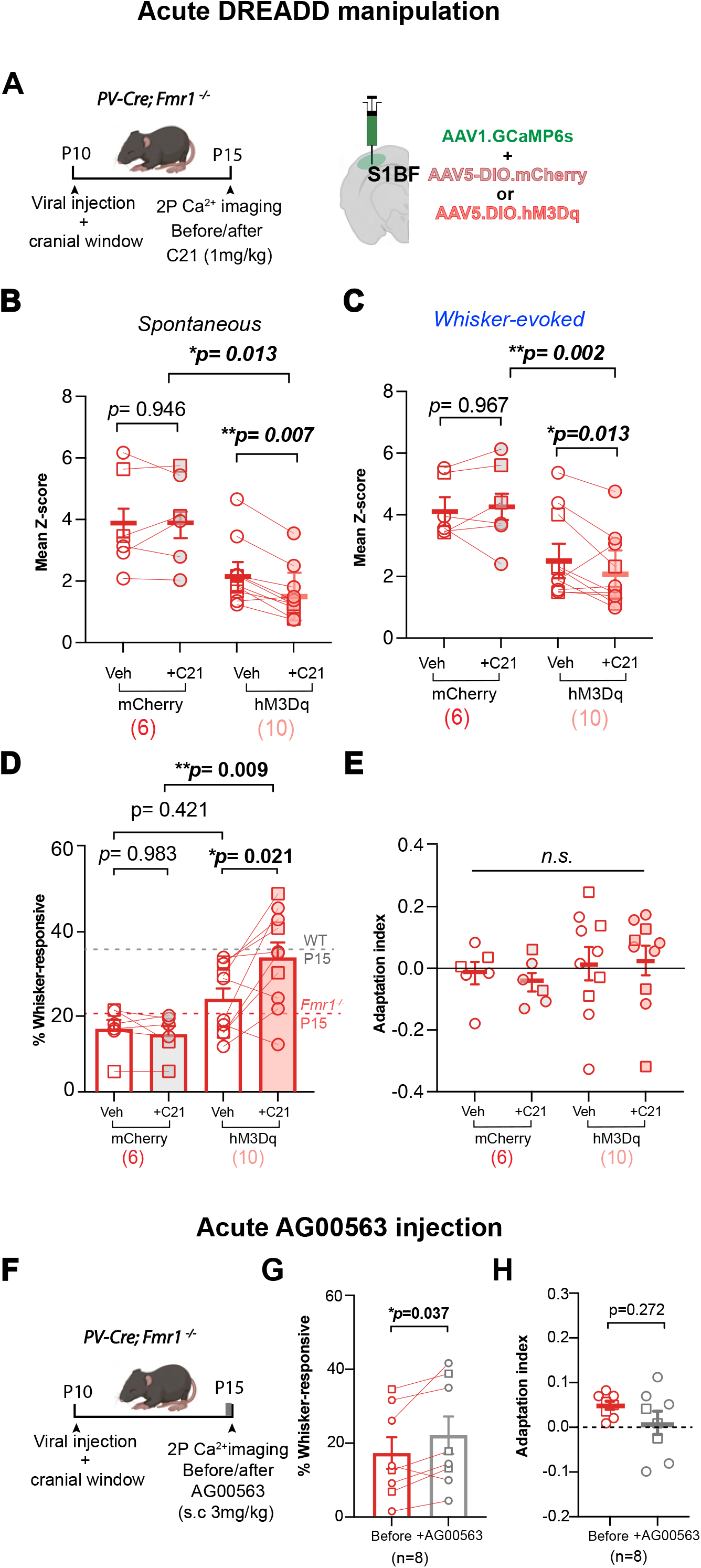
Acute activation of PV-INs in *Fmr1*^−/−^ mice at P15 increases the percentage of whisker responsive Pyr cells, but does not change their adaptation. a. Experimental design for calcium imaging recordings in Pyr cells after acute chemogenetic activation of PV-INs at P15 with the hM3Dq DREADD agonist C21 (injected ∼30 min prior to recording). b. Mean z-score for spontaneous activity of L2/3 Pyr cells is reduced upon C21 injection in *Fmr1*^−/−^- hM3Dq mice but not in *Fmr1*^−/−^ mCherry controls (before/after C21: 3.9 ± 0.7 vs. 4.0 ± 0.6 in mCherry group, p=0.946; and 2.3 ± 0.3 vs. 1.6 ± 0.3, p=0.007 in hM3Dq group; n=6 and 10 mice, respectively; two-way ANOVA with post-hoc Tukey). c. Same as panel B but for whisker-evoked activity (before/after C21: 4.2 ± 0.4 vs. 4.3 ± 0.6 in mCherry group, p=0.967; and 2.7 ± 0.4 vs. 2.2 ± 0.4 in hM3Dq group, p=0.013; two-way ANOVA with post-hoc Tukey). d. The percentage of whisker-responsive Pyr cells at P15 was significantly higher in upon C21 injection in *Fmr1*^−/−^-hM3Dq mice but not in *Fmr1*^−/−^ mCherry controls. Dashed black and red horizontal lines indicate the mean percentage of whisker-responsive Pyr neurons for WT and *Fmr1*^−/−^ mice, respectively, from our previous study (C. X. He et al., 2017). (hM3Dq group before/after C21: 23.5 ± 2.8% vs. 33.9 ± 3.9%; p=0.021; and mCherry group before/after C21: 15. 9 ± 2.3% vs. 14.5 ± 2.1%; p=0.983; two-way *ANOVA* with post-hoc Tukey). e. Neuronal adaptation was not affected by the DREADD manipulation (mCherry group before/after C21: -0.02 ± 0.09 vs. -0.05 ± 0.03, hM3Dq group: 0.02 ± 0.05 vs. 0.03 ± 0.05, p>0.99; two-way *ANOVA* with post-hoc Tukey). f. Experimental design for the acute administration of AG00563 (3 mg/kg, s.c.) and calcium imaging at P15, before and 30 min after injection. g. The percentage of whisker-responsive Pyr cells in *Fmr1*^−/−^ mice was significantly higher after AG00563 injection compared to baseline (17.1 ± 4.3% baseline vs. 21.9 ± 5.1% ∼30-40 min after AG00563, p=0.033; paired t-test, n=8 mice). h. The neuronal adaptation index of Pyr cells was not changed by AG00563 (0.05 ± 0.01 baseline vs. 0.01 ± 0.03 after AG00563, p=0.033; paired t-test, n=8 mice).

**Supplementary data Fig.10:**
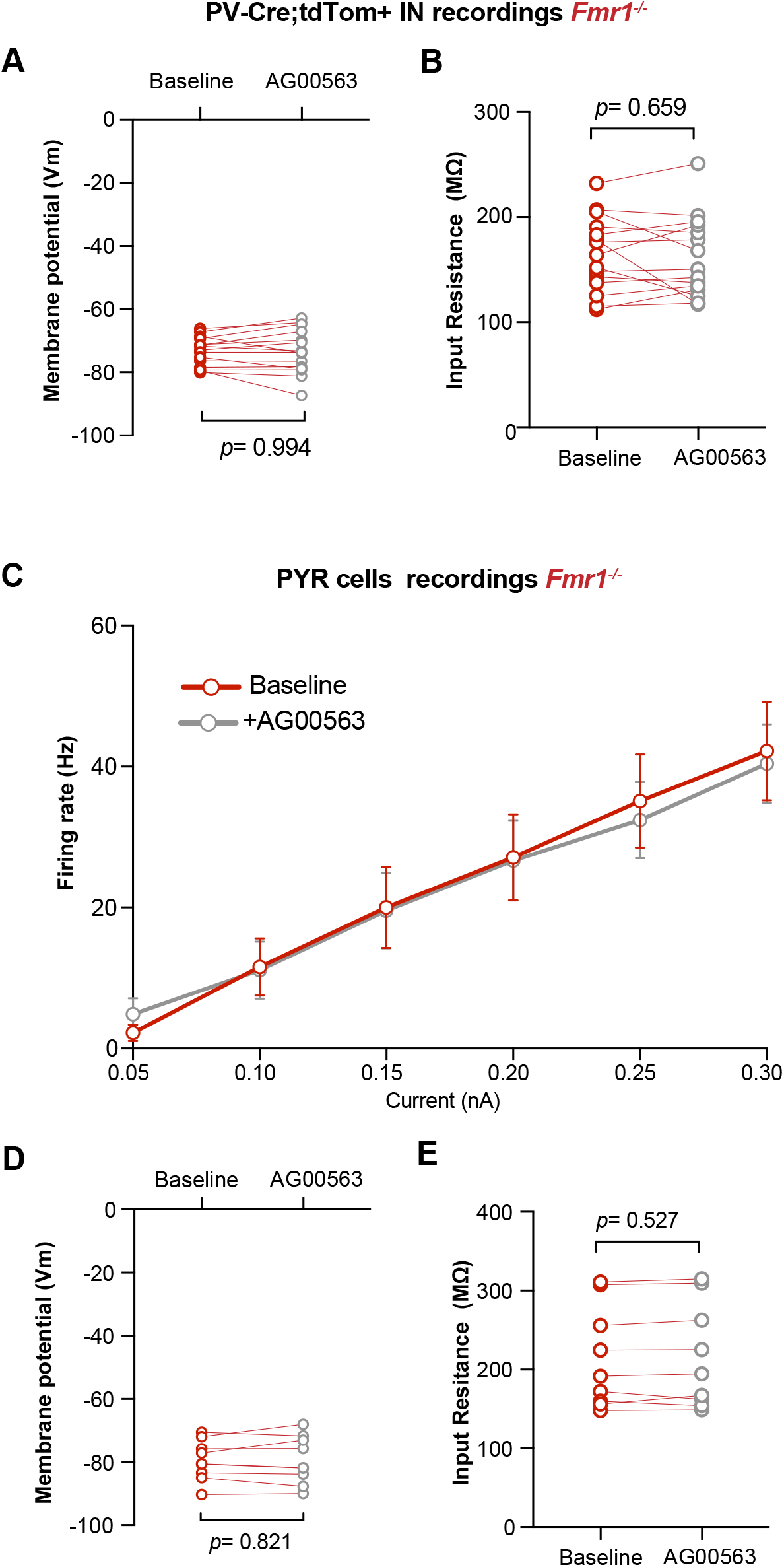
Intrinsic properties of PV-INs and Pyr cells are unchanged by AG00563. a. Resting membrane potential (Vm) of PV-INs is unchanged by bath application of AG00563 during current clamp recordings of *PV*-tdTom^+^ cells (-73.4 ± 1.2 mV vs. -73.2 ± 1.8 mV, p= 0.805, paired t-test, n=15 cells from 6 *Fmr1^−/−^*mice at P15-16). b. Input resistance (Rm) of PV-INs is unchanged by AG00563 (164.6 ± 9.2 MΩ vs. 161.2 ± 10.1 MΩ, p= 0.608, paired *t*-test). c. Cumulative input-output curves during baseline (red) or bath application of AG00563 (gray) (n=9 Pyr cells from 6 *Pv*-Cre^+/−^;tdTom^+/−^;*Fmr1*^−/−^mice, two-way RM ANOVA). d. V_m_ of Pyr cells is unchanged by AG00563 (-79.5 ± 2.1 mV vs -79.3 ± 2.5 mV, p=0.805, paired t-test). e. R_m_ of Pyr cells is unchanged by AG00563 (214.0 ± 21.4 MΩ vs. 215.4 ± 22.0 MΩ, p=0.608, paired *t*-test).

